# Estimating the rate of sexual reproduction and the inbreeding rate in *Leishmania*

**DOI:** 10.1101/2024.10.15.618469

**Authors:** Andrew G. Nicoll, Hannah Kilford, Cooper Alastair Grace, João Luís Reis-Cunha, Daniel C. Jeffares, George W. A. Constable

## Abstract

Many eukaryotic species undergo sexual reproduction facultatively, either in response to stress or in particular environments. For the remainder of their life cycle they reproduce by mitosis. This facultative sex may be rare, which alters features of their evolutionary dynamics. Both the frequency of sex and the degree of inbreeding have been challenging to estimate from genome-scale polymorphism data. Here, we describe a method to estimate both these parameters based on the Moran model of heterozygous sites. We apply this method to population genomic data from *Leishmania* parasitic protozoans from the *Leishmania donovani* species complex, showing that these parasites undergo sexual reproduction approximately once per 10,000 generations and that populations vary considerably in the extent of their inbreeding.

---

**I**n this analysis, we examine data from several populations of the *Leishmania* Trypanosomatids. Trypanosomatid parasites are a group of protozoans that cause devastating diseases, imposing severe health and economic burdens primarily upon developing countries (Burza *et al*. 2018; Kennedy 2019; Horn 2022). Among them, African trypanosomiasis, American trypanosomiasis and leishmaniasis, caused respectively by *Trypanosoma brucei*; *Trypanosoma cruzi* and species from the *Leishmania* genus are Neglected Tropical diseases (NTDs), with more than one billion people living at risk of infection. These diseases are a part of the WHO NTDs elimination road map for 2021-2030 (WHO/UCN/NTD/2020.01) (Horn 2022).

Trypanosomatids have digenetic life cycles, with developmentally distinct stages in the mammalian host and insect vectors where they sexually reproduce. Insect vectors vary; *Leishmania* species are transmitted by various species of Phlebotomine sand flies (Ready 2013), tsetse Flies (*Glossina*) as vectors of *Trypanosoma brucei* the causal agents of African trypanosomiasis and animal trypanosomiasis (Wamwiri and Changasi 2016), and Triatomine bugs for *Trypanosoma cruzi*, the agent of Chagas disease (De Oliveira *et al*. 2018; Carbajal-De-La-Fuente *et al*. 2022).

Historically, and there has been debate about whether Trypanosomatids are predominantly clonal (Gutiérrez-Corbo *et al*. 2021), but current evidence from laboratory-generated crosses and population genetics data show that many species of *Leishmania, Trypanosoma brucei* and *Trypanosoma cruzi* all undergo sexual recombination and some degree of outbreeding (Rougeron *et al*. 2009; Rogers *et al*. 2014; Messenger and Miles 2015; Kay *et al*. 2022; De Los Santos *et al*. 2022; Ferreira and Sacks 2022). Genetic exchange has been also demonstrated in the non-pathogenic Trypanosomatid species *Crithidia bombi*, occurring in the natural host, the bumblebee (Schmid-Hempel *et al*. 2011). At present, haploid gametes and mating has only been observed in *Trypanosoma brucei* (Peacock *et al*. 2014).

Indications are that the extent of outbreeding differs between clades of Trypanosomatids, species, and populations (Rougeron *et al*. 2009; Rogers *et al*. 2014; Messenger and Miles 2015; Kay *et al*. 2022; De Los Santos *et al*. 2022). For example, some populations of *T. brucei gambiense* in Africa appear to be strictly asexual (Weir *et al*. 2016). Here, we focus mainly on understanding *Leishmania* populations, although the methods we develop will have use in any diploid species with rare sex, including protozoans *Trypanosoma, Giardia, Trichomonas*, as well as many species of fungi and algae.

In *Leishmania*, genetic exchange has been shown to occur within sand fly vectors (Akopyants *et al*. 2009). *Leishmania* are capable of self fertilisation, within-species outbreeding and between species hybridisation (Gutiérrez-Corbo *et al*. 2021). For outbreeding to take place, two genotypes must be present within the insect vector. There are various ways this can occur. While multi-genotype infections do occur they appear to be rare (Cupolillo *et al*. 2020; Franssen *et al*. 2021a). However, the sand fly vectors are likely to feed on more than one host, since second blood meals are known to promote *Leishmania* propagation (Serafim *et al*. 2018), and infection of sand flies by *Leishmania* causes an increase in vector biting persistence, promoting multiple host feeding (Rogers and Bates 2007). *Leishmania* appear to have adapted to undergo meiosis in response to blood meals, since 2 Estimating the rate of sexual reproduction in *Leishmania* the formation of parasite structures that promote mating is enhanced when parasites are exposed to a second blood meal containing pre-immune IgMn antibodies (Serafim *et al*. 2023). Unlike Apicomplexan species such as *Plasmodium*, sexual cycles do not appear to be an obligatory stage for development of *Leishmania* parasites through insect vectors (Gibson 2021; Catta-Preta *et al*. 2023).

Although direct observations of cell fusion, nuclear fusion, haploid gametic cells have not yet been made, experimental crosses have now been generated in a variety of *Leishmania* species (Ferreira and Sacks 2022). While *in vitro* methods are available, selection of couple-drug resistant F1 offspring is most effective from crosses performed in sand flies (Ferreira and Sacks 2022). Studies of experimental crosses generate offspring with heterozygosity expected from equal contributions of haplotypes from parental strains, and are consistent with a meiosis-like process (Inbar *et al*. 2019; Louradour *et al*. 2020; Shaik *et al*. 2021; Serafim *et al*. 2023; Sádlová *et al*. 2024).

Population genomic data analysis of various *Leishmania* species and populations indicate that inbreeding, within-species outbreeding and between species hybridisation all occur in natural conditions (Belli *et al*. 1994; Delgado *et al*. 1997; Rogers *et al*. 2014; Schwabl *et al*. 2019; Cotton *et al*. 2020; Van den Broeck *et al*. 2020; Kato *et al*. 2021; Lypaczewski and Matlashewski 2021; Glans *et al*. 2021; De Los Santos *et al*. 2022; Grace *et al*. 2022; Pilling *et al*. 2023; Hadermann *et al*. 2023). The extent of inbreeding appears to differ between species and populations. Interspecies hybrids appear to common, for example between *L. aethiopica* and *L. donovani* in East Africa (Hadermann *et al*. 2023), *L. donovani* - *L. major* and *L. donovani* - *L. tropica* hybrids in Sri Lanka (Lypaczewski and Matlashewski 2021). *L. peruviana* - *L. braziliensis* Peru (Van den Broeck *et al*. 2020), and a variety of interspecies hybrids within the Viannia clade (*L. braziliensis, L. peruviana, L. lainsoni, L. guyanensis* and *L. shawi*) have also been discovered in Peru (Kato *et al*. 2021).

*Leishmania* and some other Trypanosomatids have some unusual genome biology that is likely to affect the extent of heterozygosity, and therefore estimates of inbreeding. The somy of chromosomes varies dynamically within populations (Black *et al*. 2023), and one chromosome is typically tetraploid (Reis-Cunha *et al*. 2024a). Full genome duplications to form triploids and tetraploids are also found within *Leishmania* populations (Van den Broeck *et al*. 2023; Hadermann *et al*. 2023). Chromosome copy numbers are subject to both genetic drift and selection, in a process called ‘haplotype selection’ (Dumetz *et al*. 2017; Prieto Barja *et al*. 2017; Bussotti *et al*. 2023). In experimental conditions, the copy numbers of some chromosomes varies predictably between insect and mammalian stages (Dumetz *et al*. 2017). The dynamics of chromosome duplication and loss may result in complete loss of heterozygosity within chromosomes, for example where an Aa haplotype pair duplicates the A haplotype to AAa, followed by loss of a, leaving two (homozygous) AA haplotypes. Consistent with the expectation from this process, extensive loss of heterozygosity (LOH) has been observed in some populations (Hadermann *et al*. 2023; Glans *et al*. 2021).

Since *Leishmania* species undergo meiosis in only in sand flies, the number of cell divisions in the insect and mammalian stages have an influence on the frequency of sex. There is likely to be a strong population bottleneck when sand flies receive *Leishmania* parasites from mammalian hosts. Laboratory studies show that sand flies ingest a median of 50-80 parasites after feeding on infected hamsters (Serafim *et al*. 2018). Using a different experimental system it has been estimated that sand flies transmit between 600-1000 parasites, with rare cases of 100,000 parasites transmitted (Kimblin *et al*. 2008). By contrast, patients with severe visceral leishmaniasis can harbour on the order of 10^7^ parasites in the blood and perhaps as many as 10^11^ parasites in the bone marrow (Verma *et al*. 2010). Therefore, the majority of *Leishmania* life cycle will be within the mammalian stage and the majority of cell divisions will be mitotic divisions in mammals.

The rate of sexual (as opposed to clonal) reproduction in natural populations is an important evolutionary quantity, particularly in pathogenic species (de Meeûs *et al*. 2007; Halkett *et al*. 2005) such as *Leishmania*. Despite this, it has proved surprisingly difficult to estimate this parameter from genomic data (Stoeckel and Masson 2014a; Ali *et al*. 2016; Ennos and Hu 2019). Methods to detect clonal reproductive life cycles include analysis of population structure, clonal diversity (i.e. the detection of multiple clones), linkage disequilibrium (LD) and finally genetic diversity (including allelic diversity and F-statistics) (Halkett *et al*. 2005). Amongst these use of LD is perhaps the most common (Tsai *et al*. 2008a). Increased levels of LD in a population are correlated with increased rates of clonal reproduction (Hill and Robertson 1968; Smith *et al*. 1993). The mean correlation coefficient of genetic distance between unordered alleles, 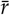,(a measure of linkage disequilibrium), can be used to infer rates of sex, however this requires large sample sizes (Hill 1981; Waples 1989). Alternatively, rates of decay of LD with distance along the genome can be used to estimate rates of sexual reproduction (Talas and McDonald 2015; Nieuwenhuis and James 2016). However these analyses require recombination rates to have been determined in advance from laboratory studies (Ennos and Hu 2019). Finally, while methods have been developed that allow quantification of rates of asexual vs sexual reproduction in populations genotyped at two time steps (Ali *et al*. 2016), these come with the downside of being incompatible with the one-time step sampling strategy of most population genetic data. As *Leishmania* population genomic surveys require isolation of parasites from patients they are seldom collected more than once per patient, rendering such time sampling methods impractical.

Recently, the use of higher-order F-statistics has been further investigated as an approach for estimating the rate of sexual reproduction in species with rare facultative sexual reproduction in diploids (Stoeckel and Masson 2014a). Wright’s inbreeding coefficient, *F*_IS_, measures departures from Hardy-Weinberg equilibrium (Wright 1931). The rate of relaxation to this equilibrium is dependent on the rate of sexual reproduction (Reichel *et al*. 2016), with frequent sex constraining populations more tightly around values of *F*_IS_ expected under Mendelian segregation. Therefore, although the mean value of *F*_IS_ may be independent of the rate of sex, it has long been known that increased variance of *F*_IS_ is correlated with less sexual reproduction (Balloux *et al*. 2003; Allen and Lynch 2012). Stoeckel and Masson (2014a) proposed using this property, along with the higher-order moments of *F*_IS_, to estimate the rate of sexual reproduction in organisms with rare sex. Indeed, more recent work applying supervised machine learning to simulated population genetic data reveals the importance of the distribution of *F*_IS_ as a means to estimate the rate of sex in highly clonal systems (with rates of clonality exceeding ≈ 95%) (Stoeckel *et al*. 2021).

In this paper we take an analogous approach to previous studies that have used departures from Hardy-Weinberg equilibrium to estimate the rate of sexual reproduction in faculatatively sexual diploids (Stoeckel and Masson 2014a; Stoeckel *et al*. 2021). However, rather than fitting the distribution of Wright’s inbreeding coefficient, *F*_IS_, we utilise the genotype distribution in a biallelic single locus model. In general, numerically obtaining such a genotype distribution is computationally challenging due to the high dimensionality of the genotype state space for realistic effective population sizes (Reichel *et al*. 2015). However, we will show mathematically that a compound parameter emerges; the population size multiplied by the probability of sex per reproductive event, or analogously the number of sexual events per generation, as has been identified in earlier studies (Ennos and Hu 2019; Hartfield 2016a). This parameter allows us to approximate the genotype distribution and make computational progress. In addition, we also incorporate an inbreeding rate parameter to our model due to its potential importance in *Leishmania* reproduction, as described above. Co-fitting both the inbreeding rate and the rate of sex parameters is important due to their joint affect on the genotype distribution; in effect inbreeding increases the mean of *F*_IS_ while decreasing the rate of sex increases the variance of *F*_IS_ (Rousset 2002).

The paper is organised as follows. In Materials and methods, we first introduce the *Leishmania* data sets that we will use in Data. We then introduce the mathematical model that we will fit this data to in Mathematical Modeling. In Parameter Fitting we describe the parameter fitting technique, including an approximation for dealing with large genotype spaces. Our Results section begins with an overview of the Model behaviour, before illustrating the success of the fitting approach with simulated data in Robustness checks on fitting. Finally we provide Estimates for rate of inbreeding and rate of sexual reproduction across geographic samples for several populations of the *Leishmania donovani* species complex.

## Materials and methods

### Data

SNP genotype data were obtained from Grace *et al*. (2021), using methods described therein. Populations were selected using Admixture Alexander and Lange (2011), with K=9 populations. From the population EA1, we excluded four samples that appeared to be multiclonal infections, as described in Franssen *et al*. (2021b). European Nucleotide Archive accessions for all samples used in this study are provided in [Supplementary Table 1]. The effective population size *N*_*e*_ for all populations was estimated based on the estimator *θ* = 4*N*_*e*_*µ*, using nucleotide diversity *π* as an estimator of *θ*. Nucleotide diversity was estimated using VCFtools v.0.1.15 Danecek *et al*. (2011). As no experimentally-determined mutation rate (*µ*) is available, we used an estimate of *µ* = 1.99 × 10^−9^, that was derived from comparative data from a range of eukaryotic species, using the established relationship between genome size and mutation rate Rogers *et al*. (2014).

**Table 1.** Datasets organised by geographical location including the number of samples, *N*_*s*_ and the number of analysed loci after filtering, *M*. Wright’s inbreeding coefficent, *F*_IS_ is also provided for each dataset, together with an estimate of the inbreeding parameter obtained from a deterministic analysis that neglects facultative sex, *β*(*F*_IS_) (see Eq. (13)). Results obtained from co-fitting the inbreeding parameter (i.e. accounting for facultative sex) show much higher levels of inbreeding 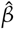.The best-fit number of sexual events per generation, *N*_*f it*_*σ*_*f it*_, are consistent across samples. The inferred best-fit rate of sexual reproduction, 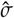,shows greater variation across samples, driven by variation in *N*_*e*_ across samples. Standard deviation errors on the inbreeding parameter, 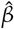,and rate of sex parameter, 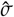,are obtained using the Fisher information index of the maximum of the likelihood surface (see Supplementary Information). Note that this is evidently an underestimate of the true uncertainty, resulting from our simplifying assumption of no genetic linkage between sites.

| Data set | $N_s$ | $N_{fit}$ | $M$ | $F_{IS}$ | $\beta(F_{IS})$ | $\hat{\beta}$ | $N_{fit}\sigma_{fit}$ | $N_e$ | $\hat{\sigma}_e$ |
| --- | --- | --- | --- | --- | --- | --- | --- | --- | --- |
| EA1 | 37 | 200 | 26,165 | 0.133 | 0.235 | $0.44 \pm (4.3 \times 10^{-3})$ | $5 \pm (4.2 \times 10^{-2})$ | 53,266 | $9.38 \times 10^{-5}$ |
| EA2 | 18 | 200 | 10,211 | 0.688 | 0.815 | $0.99 \pm (1.8 \times 10^{-3})$ | $11.5 \pm (1.5 \times 10^{-1})$ | 10,930 | $1.1 \times 10^{-3}$ |
| ISC1 | 211 | 250 | 355 | 0.492 | 0.659 | $0.93 \pm (9.9 \times 10^{-3})$ | $5.1 \pm (3.1 \times 10^{-1})$ | 10,553 | $4.9 \times 10^{-4}$ |
| ISC2 | 15 | 200 | 5,961 | 0.633 | 0.776 | $1 \pm (1.0 \times 10^{-3})$ | $5.8 \pm (9.8 \times 10^{-2})$ | 791 | $7.3 \times 10^{-3}$ |
| Brazil | 127 | 200 | 2,123 | -0.144 | -0.33 | $0.75 \pm (2.0 \times 10^{-2})$ | $1.8 \pm (4.8 \times 10^{-2})$ | 3,518 | $5.1 \times 10^{-4}$ |

### Mathematical Modeling

We consider a single locus diploid Moran-type model of fixed population size *N* (i.e. generations are overlapping and birth-death events are coupled), with alleles A and a (Moran 1958; Ewens 2004). We are particularly interested in the rare-sex regime, where departures from Hardy-Weinberg equilibrium are large; indeed, it is these departures that allow us to estimate the rate of sexual reproduction using segregation data alone. We therefore need to capture the specific abundances of AA, Aa, and aa genotypes, with abundances denoted by the variables *n*_*AA*_, *n*_*Aa*_, and *n*_*aa*_ = *N* − *n*_*AA*_ − *n*_*Aa*_ (Hössjer and Tyvand 2016; Constable and McKane 2018). Sexual reproduction occurs at a probability rate *σ*, while asexual (clonal) reproduction occurs at a probability rate (1 − *σ*). We further assume that the genotypes are selectively neutral with respect to one another.

As we have addressed, sexual reproduction in *Leishmania* is assumed to occur within the sand fly vector. Low genetic diversity in the sand-fly’s blood meal can therefore lead to high rates of inbreeding. We account for this by introducing an inbreeding parameter *β*; once an individual is selected for sexual reproduction (with probability *σ*), with probability *β* that individual mates with a partner of its own genotype, while with probability (1 − *β*) it mates with a partner selected at random from the entire population (within the insect in this case). With *β* = 0 (i.e. no inbreeding), the model then amounts to a continuous-time version of the model considered by Stoeckel and Masson (2014b), albeit with a clonality parameter *c* replaced with a sexual parameter *σ* such that *σ* = (1− *c*).

The state space of the model is illustrated in Figure 1. From an interior state (away from the boundaries where one of the genotypes is extinct) there are six transitions taking the population from ***n*** = (*n*_*AA*_, *n*_*aa*_) to neighbouring states. The rates at which these transitions occur are denoted 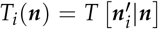. The full list of these rates in given in the Supplementary Information S2, however here we use *T*_5_(***n***) as an illustrative example. Assuming Mendelian inheritance, we have

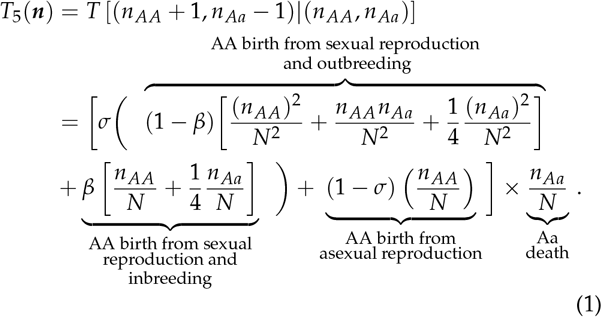

where we have allowed for selfing within the population.

**Figure 1.**
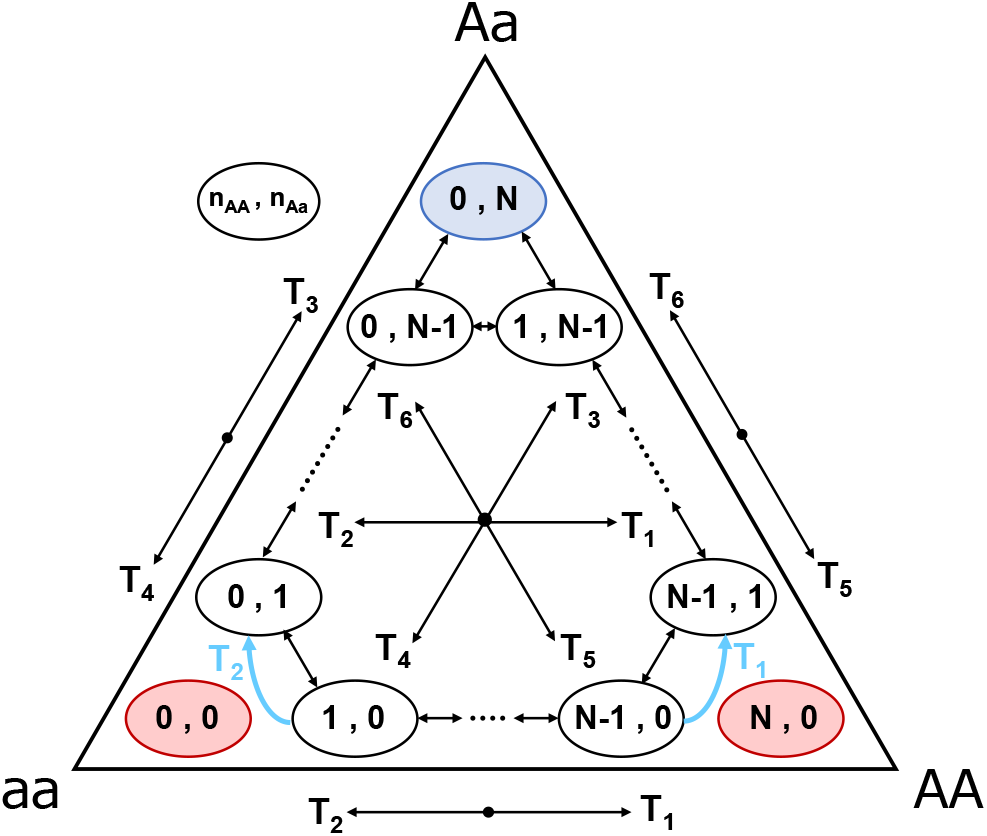
State space and transitions in the mathematical model (see, for instance, Eq. (1)). Each state corresponds to a unique combination of specific abundances of the AA and Aa genotypes, denoted (*n*_*AA*_, *n*_*Aa*_). Transitions are represented by arrows between states, with the six transition rates, denoted *T*_*i*_, corresponding to a fixed direction as seen in the centre. Red states correspond to absorbing states in which one of the alleles (A or a) has become extinct. Blue state represents the state of maximum heterozygosity. Blue rewired transitions (*T*_2_ left, and *T*_1_ right) represent the introduction of a new mutant to the population after the extinction of one allele (see Mathematical Modeling).

States ***n*** = (*N*, 0) and ***n*** = (0, 0) (represented in red in Figure 1) are absorbing states of the model, from which the population cannot escape in the absence of mutation. However, as our data is filtered to remove such non-segregating sites, we rewire our transitions such that those involving the deaths of the last remaining homozygote instead take the system to a state with a single heterozygote (see blue transitions in Figure 1). This rewiring can be interpreted as the instantaneous introduction of a new mutant upon extinction of either allele and comes with the advantage of not having to specify a mutation rate for the model. Note that this also implicitly assumes at least some low level of sexual reproduction (i.e. *σ* > 0) such that transitions from the state ***n*** = (*n*_*AA*_, *n*_*Aa*_) = (0, *N*) are dominated by segregation (i.e. *σ*≫ *µ*).

The model dynamics for the probability of being in state ***n*** at time *t, P*_***n***_ (*t*), are then specified by Kolmogorov’s forward equation (also known as the the master equation), which is denoted (Ewens 2004; Gardiner 2009)

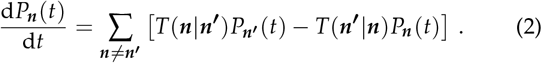

This equation simply states that the probability that the system is in state ***n*** increases with the probability of entering from state 4 Estimating the rate of sexual reproduction in *Leishmania* ***n***^***′***^ and decreases with the probability of being in state ***n*** and leaving to enter state ***n***^***′***^. This equation is expressed in terms of our probability transitions rates *T*_*k*_ (***n***) in the Supplementary Information S2. Of particular interest to us is the stationary distribution of Eq. (2), to which *P*_***n***_ (*t*) equilibriates at long times. This distribution, which we denote by 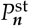, is obtained by setting d*P* (*t*)/d*t* = 0 in Eq. (2) and solving for 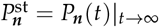,

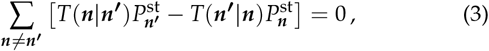

such that the distribution is time-independent at equilibrium. In the following sections, we will solve these equations (which amount to an eigenvalue-eigenvector problem) numerically (Gardiner 2009; Ewens 2004). However for building intuition, it is useful to consider applying a diffusion approximation to the model (Watterson 1964; Ewens 2004).

Assuming that *N* is large, we introduce the approximately continuous vector of genotype frequencies ***x*** = ***n***/*N*. Then conducting a Taylor expansion of Eq. (2) in the small parameter *N*^−1^ and keeping only the two leading order terms we obtain the Fokker-Planck equation for the model, an advection-diffusion equation for the continuous probability distribution *p*(***x***, *t*), which is itself analogous to *P*_***n***_ (*t*) (Risken 1989). The resulting equation can be expressed as

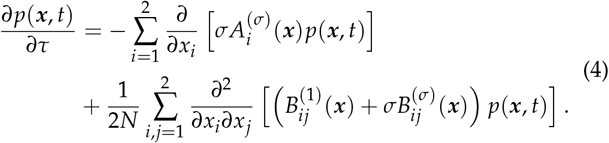

where time is measured in units of generations (i.e. *τ* = *Nt*) and we have taken care to separate out terms of different orders in *N* and the probability of sexual reproduction, *σ*, and truncated terms of order *N*^−2^. These terms are given in full in Supplementary Information S2.

In an analogous manner to the way the discrete stationary probability distribution 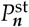 can be calculated from Eq. (2), the continuous stationary probability distribution *p*^st^ (***x***) = *p*(***x***, *t*) |_*t →*∞_ (which approximates 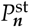) can be calculated from Eq. (4). Of particular interest to this current study is the case when sexual reproduction is rare; if *σ* is of the order *N*^−1^ (i.e. there are just a few sexual events per generation). In this case the terms 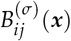 in Eq. (4) can be neglected (since they are of order *N*^−2^), and the continuous stationary probability distribution is given by the solution to

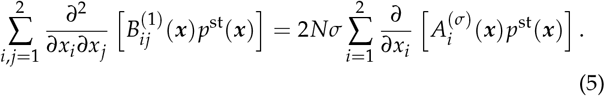

Importantly, we now see that *σN*, which can be interpreted as the expected number of sexual events per generation, now appears as a compound parameter. This implies that when *N* is large (such that the diffusion approximation in Eq. (4) remains valid and *σ* is small (such that the approximation in Eq. (5) remains valid, the stationary probability distribution remains unchanged so long as *σN* is held constant. This echos results obtained in Ennos and Hu (2019). We shall make use of this insight in the following section.

Finally, we note that if we take the limit of infinite population size (*N* → ∞), Eq. (4) simplifies to the following set of ordinary differential equations (Risken 1989; Gardiner 2009),

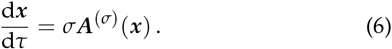

where ***A***^(*σ*)^(***x***) retains the same form as in Eq. (4) (see also Supplementary Information S2). Here we see that in the deterministic limit (i.e. in the absence of genetic drift) the timescale of the dynamics are entirely governed by the rate of sexual reproduction *σ*, with low rates of sex implying a slow approach to equilibrium (Reichel *et al*. 2016).

### Parameter Fitting

We begin by considering an empirical population with an effective population size *N*_*e*_, and assume that the population is at equilibrium such that we can fit data obtained from this population to the stationary probability distribution 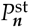 for the model (see Mathematical Modeling). In practice genotype data is sampled from a small number of isolates from 18 to 211 in our case, see Table 1; we denote this number of samples by *N*_*s*_. Genotype data is filtered to remove loci with a minor allele frequency less than 5%, as well as loci for which the number of segregating alleles is not equal to two (see Data). We denote the number of remaining segregating loci by *M*. For simplicity, we will assume no genetic linkage between these loci.

In order to reflect the sampling described above, we introduce a sampled distribution

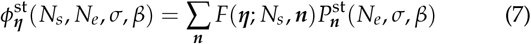

where *F*(***η***; *N*_*s*_, ***n***) is a multinomial distribution that gives the probability of sampling ***η*** = (*η*_*AA*_, *η*_*Aa*_) homozygotes and heterozygotes at a given loci following following *N*_*s*_ samples of the population, and we have introduced the parameter dependence of each stationary distribution for clarity. Note that this implicitly assumes that *N*_*e*_≫*N*_*s*_, such that effects of resampling individuals can be ignored. In order to account for the data filtering described above in our mathematical model, we further renormalise the sampled stationary distribution:

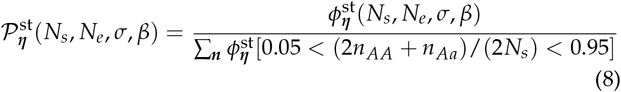

where the square brackets denote an Iverson bracket and we have introduced a functional dependence on *N, σ*, and *β* to emphasise the dependence of the stationary distribution on these model parameters.

Assume now that at the *i*^*th*^ locus the empirically sampled abundance of AA and Aa types in the population is measured to be ***m***_*i*_ = (*m*_*AA*_, *m*_*Aa*_)_*i*_ for *i* = 1, …, *M*, where *N*_*s*_≥ *m*_*AA i*_ + *m*_*Aa i*_ for all *i*. The likelihood of this observation in our mathematical model for given parameters is simply 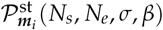. Therefore the log-likelihood of observing all *M* segregating sites is

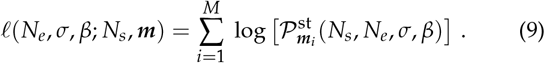

where we have for simplicity assumed no genetic linkage, a common simplification. The best fit parameters for *N*_*e*_, *σ*, and *β*, denoted 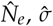,and 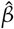are then obtained by maximising the likelihood function such that

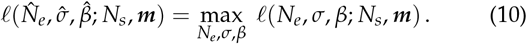

While the above direct approach allows us to estimate the population size, rate of sexual reproduction and inbreeding rate, it has computational drawbacks. Obtaining 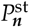 in Eq. (8) requires solving (*N*_*e*_ + 1)(*N*_*e*_ + 2)/2 difference equations, which can be prohibitively expensive computationally for application to *Leishmania*, which has often has effective population sizes of order 10^4^ (Table 1). However, a computationally cheaper approach can be 6 Estimating the rate of sexual reproduction in *Leishmania* obtained by exploiting the results obtained using the diffusion approximation in the previous section.

In Eq. (5) we saw that when the population size was large and sex was rare (such that *σN* ≈ 𝒪 (1)), the stationary distribution for the population was approximately unchanged when *σN* was held constant. A modified (and computationally cheaper) version of the fitting process described above can therefore be derived in which the compound parameter *N*_fit_ *σ*_fit_ can be estimated with a fixed *N*_fit_;

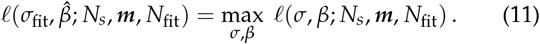

For *N*_*e*_≫ *N*_fit_≫ 1, the speed of the fitting algorithm can be increased, while the accuracy of the diffusion approximation (from which we infer the key role of the compound parameter *σN*) is maintained. Eq. (11) then allows us to estimate both the inbreeding rate, 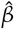,and the number of sexual events per generation, *N*_fit_ *σ*_fit_. The actual probability rate of sexual reproduction in the population, 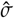,can then be inferred if we have an independent approximation of the effective population size;

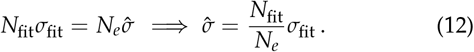

This approach is discussed in more detail in Supplementary Information S4.2. Essentially by reducing the size of the genotype state space that we need to consider, we are able to fit the data to the genotype distribution directly rather than needing to fit the distribution of *F*_IS_ (Stoeckel and Masson 2014b; Stoeckel *et al*. 2021).

### Statistical analysis

Log-likelihood surfaces are obtained for the fit of the datasets to the model in the *Nσ* − *β* plane according to Eq. (9), as illustrated in Section S6 of the Supplementary Information. The best fit parameters for the datasets are then obtained by locating the maximum of the log-likelihood surfaces (see Eq. (10)). In order to obtain an estimate of the uncertainty associated with these maximum likelihood estimates, we first compute the Fisher information matrix. The Fisher information matrix measures the curvature of the likelihood surface (it is the Hessian matrix of the likelihood surface). Since our likelihood surface is obtained numerically, we calculate this curvature to first-order numerically. The covariance matrix (estimating the uncertainty in the fitted parameters 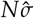 and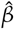) is then be calculated as the inverse of the Fisher information matrix (Kagan and Landsman 1999; Mielke and Schwabe 2010). This covariance matrix is illustrated in Figs S7-S11 of the Supplementary Information. The uncertainties in Table 1 are then the standard deviations obtained from this covariance matrix. The analysis assumes for simplicity that the samples of genotype frequencies at each locus are statistically independent (i.e. that all loci are unlinked) resulting in an under estimate of the true uncertainty in estimates of the inbreeding rate and rate of sexual reproduction,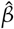 and 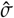 respectfully.

## Results

### Model behaviour

For orientation, we begin by discussing the behaviour of the model. In the infinite population size (deterministic) limit, the dynamics are given by Eq. (6). So long as sex occurs at a non-zero rate (*σ* > 0), the population approaches a line of fixed points (the centre manifold, or CM, see Eq. (S44)) at a rate proportional to *σ* (see red dashed lines in left panels of Fig. 2). For the case of no inbreeding (*β* = 0), this is simply the Hardy-Weinberg equilibrium. For *β* > 0, inbreeding reduces the proportion of heterozygotes at equilibrium. An analysis of the CM shows that in this infinite population size limit, *β* is related to Wright’s inbreeding coefficient, *F*_*IS*_, via

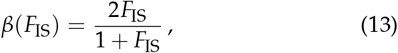

as illustrated by Pollak (1987) (see also, Supplementary Information S2.3).

**Figure 2.**
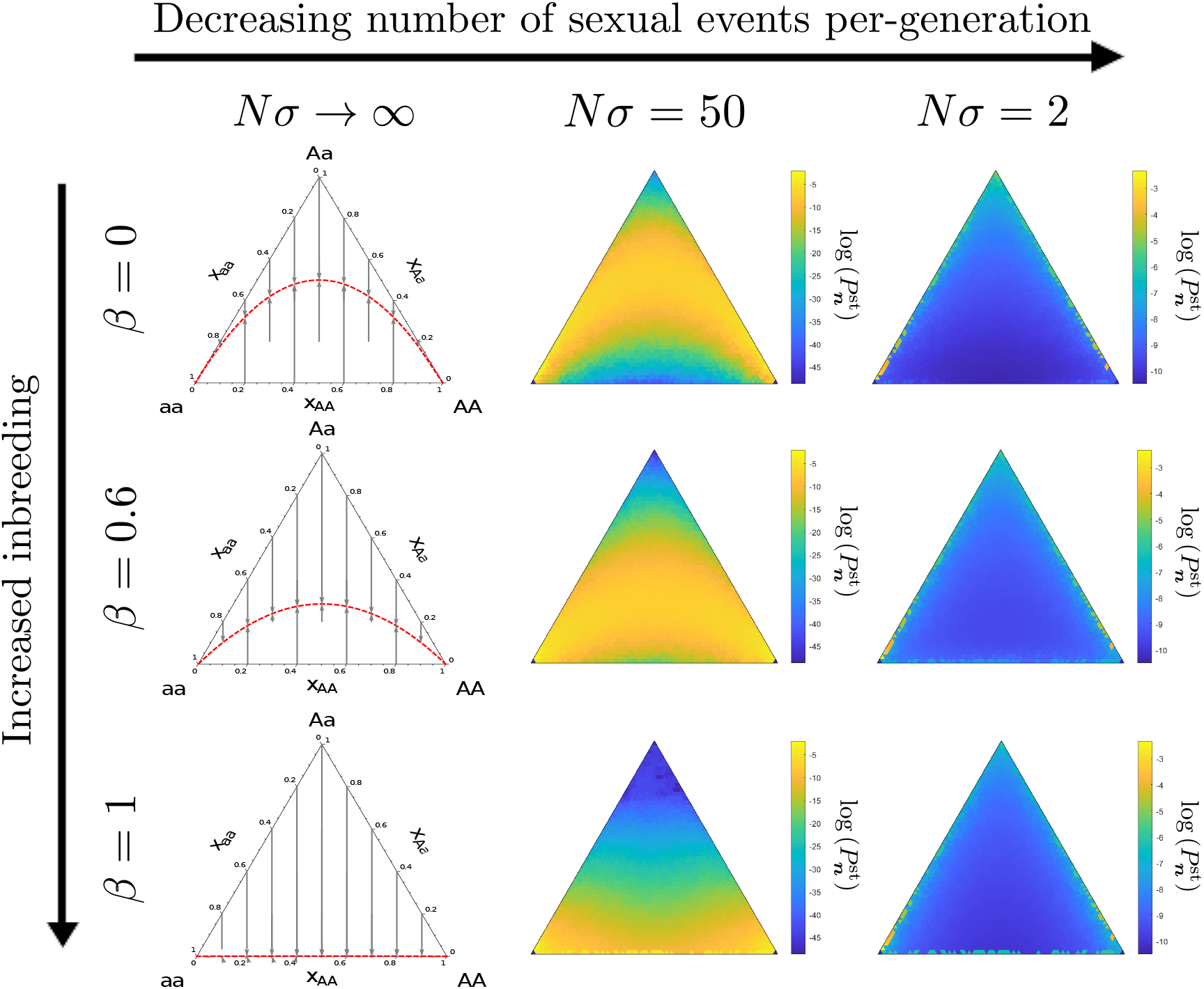
Illustrative model behaviour in various parameter regimes. Left panels: As the number of sexual events per generation, *Nσ* tend to infinity, the dynamics converge on the deterministic limit (see Eq. (6)), and trajectories approach a centre-manifold representing the Hardy-Weinberg equilibrium (red dashed lines). Central Panels: When *Nσ* is large but finite, genetic drift leads to a stationary probability distribution around the manifold predicted in the deterministic limit. Right panels: When *Nσ* is small, temporary extinctions of the heterozygoes and homozygotes become more likely, and the deterministic analysis no longer forms a useful approximation of the long-time behaviour. Nevertheless, the effect of sexual reproduction and inbreeding still leaves a signature on the resultant distribution, with the probability of low heterozygote numbers increasing with inbreeding. Parameters: In all central and right panels, *N* = 50; in central panels *σ* = 1; in right panels *σ* = 0.04. Note that apparent asymmetries in the distributions are a result of plotting (see Supplementary Information S7).

We next turn to the case of finite populations. In this case, we can numerically solve Eq. (3) for the stationary distribution 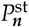to compare how the stationary probability changes as the various parameters in the model are varied (see Fig. 2, central and right panels).

When the number of sexual reproduction events per generation is large (*σN*≫1), genetic drift acts to create a distribution around the CM (see Fig. 2, central panels). For 1 > *β* and *σN* ≫ 1, this distribution is approximately Gaussian perpendicular to the CM (i.e. in the stable direction), with a mean centred on the proportional centre manifold and a variance that is proportional to 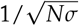 (Van kampen 2007; Stoeckel *et al*. 2021). For *β* = 1, the CM lies coincident with the boundary *n*_*Aa*_ = 0, and therefore it is clear that the distribution about the CM can no longer be symmetric, and therefore it is non-Gaussian. However the general qualitative picture remains that decreasing *Nσ* increases the variance in heterozygosity.

When the number of sexual reproduction events per generation is small (*σN* ≈ 1), the deterministic dynamics identified in the *N* →∞ limit no longer form a useful indicator of the equilibrium population dynamics (compare Fig. 2, right panels with central panels). Instead, frequent asexual reproduction leads to frequent “extinctions” of one or other of the genotypes, so that probability builds up around the boundaries (Fig. 2, right panels). The spread of the distribution away from these boundaries *increases* with *Nσ* (which increases the rate of escape from these temporary “extinction” boundaries through sexual reproduction). When inbreeding is absent (*β* = 0), we see a high concentration of probability near the state of full heterozygosity (*n*_*Aa*_ ≈ *N*), and a very low concentration of probability around states of zero heterozygosity (*n*_*Aa*_ ≈ 0), as illustrated in (Fig. 2, top-right panel). For intermediate rates of inbreeding, there is a greater concentration of probability on the interior of the state space (see Fig. 2, middle-right panel). Finally at full inbreeding (*β* = 1) there is a high probability concentration of probability at zero heterozygosity (*n*_*Aa*_ = 0), as shown in Fig. 2, bottom-right panel.

### Robustness checks on fitting

The maximum likelihood calculation in Eq. (11) depends on three fitting parameters. The first, *N*_*f it*_, relates to our calculation of the stationary distribution 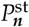,which makes the fitting approach computationally feasible. The next two parameters relate to the data; the number of segregating loci, *M* = ||***m***||, and the number of individuals sampled, *N*_*s*_. In order to explore the sensitivity of our approach to changes in these parameters, we simulate the model described by Eq. (2) using the Gillespie Algorithm Gillespie (1977). This gives us data sets with known *N* = *N*_*e*_, *β* and *σ* which we can use to test the results of our fitting approach (see Supplementary Information S4.2).

In Fig. S4 we explore what happens to accuracy of our fitted estimates as the parameter *N*_*f it*_ is reduced. We first note that the *Nσ* rule that was identified analytically in Mathematical Modeling emerges empirically from simulated data. That is, given a simulated population of actual size *N*_*sim*_ = *N*_*e*_ fitted to a sampled distribution 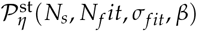 with *N*_*e*_ > *N*_*f it*_, the maximum likelihood estimate for *σ*_*f it*_ is a factor *N*_*e*_ /*N*_*f it*_ greater than than the actual simulated *σ* (see Fig. S4 (A-B)). The maximum likelihood value of the true *σ* can then be obtained using Eq. (12), as illustrated in Fig. S4 (C). We also see that when *N*_*f it*_ becomes too small (50 ⪆ *N*_*f it*_), the rate of inbreeding 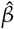 is overestimated (see Fig. S4 (D)). This effect stems from the very high level of discretization of the 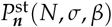 state space when *N* is small.

In Fig. S5 we illustrate the effect of reducing the number of observations of the simulated data. We find that very low numbers of observations (1000 ⪆ *M*) can lead to an over-estimate of 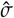 (see Fig. S5 (A)) and both an over and under-estimate of 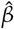 (see Fig. S5 (B)). This is equivalent to a reducing the number of segregating sites, *M*, in our data if there was no linkage between loci.

Finally in Fig. S6 we explore what happens as the number of individuals sampled from our simulated dataset, *N*_*s*_, is reduced. We find that a small number of samples (50 ⪆ *N*_*s*_) can lead to an underestimate for the rate of sexual reproduction, 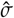 (see Fig. S6 (A)), and overestimate for the rate of inbreeding 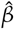 (see Fig. S6 (B)).

### Importance of co-fitting rate of inbreeding and rate of sexual reproduction under rare sex

The definition of Wright’s inbreeding coefficient, *F*_IS_, compares observed and expected heterozygosity, *H*_obs_ and *H*_exp_ respectively (). In terms of the model we are using, this takes the simple form

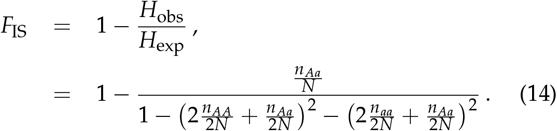

Here it is clear that any population genetic processes aside from inbreeding that causes an excess of heterozygotes can lead to an underestimate of this statistic, even driving it negative (Schwabl *et al*. 2021). Rare facultative sexual reproduction is just such a process (Hartfield 2016b). This effect can be understood intuitively by considering Figure 2, where one can see that under rare sex (right panels), the stationary probability distribution is more heavily weighted towards high heterozygote frequencies than one would expect under frequent sexual reproduction (see Figure 2, central panels). Thus the extent of inbreeding can be masked by *F*_IS_ in facultatively sexual populations, leading to an underestimate of the inbreeding rate *β* as determined by Eq. (13).

The approach described in this paper provides an alternative to using Eq. (13). By co-fitting the inbreeding parameter *β* together with *σ*_fit_ *N*_fit_ (see Eq. (11)), we obtain a more accurate estimate of *β* that is suitable for faculatatively sexual populations. We illustrate this in Supplementary Information, Fig. S6 (B), where we show using simulated data that our *β* estimates are a large improvement over those obtained from Eq. (13), which inappropriately assumes a population at deterministic equilibrium.

### Estimates for rate of inbreeding and rate of sexual reproduction across geographic samples

In Figure 3 we provide a visualization of the data for one of our datasets, the *Leishmania donovani* EA1 population from East Africa (see panel (a)), alongside the maximum likelihood fit obtained from our model (see panel (b)). In the Supplementary Information we provide the full likelihood surfaces, data visualisation and maximum-likelihood fits of the model to the data for all of our geographical datasets: EA1 (see Fig. S7), EA2 (see Fig. S8), ISC1 (see Fig. S9), ISC2 (see Fig. S10), Brazil (see Fig. S11). These results are summarised in Table 1.

**Figure 3.**
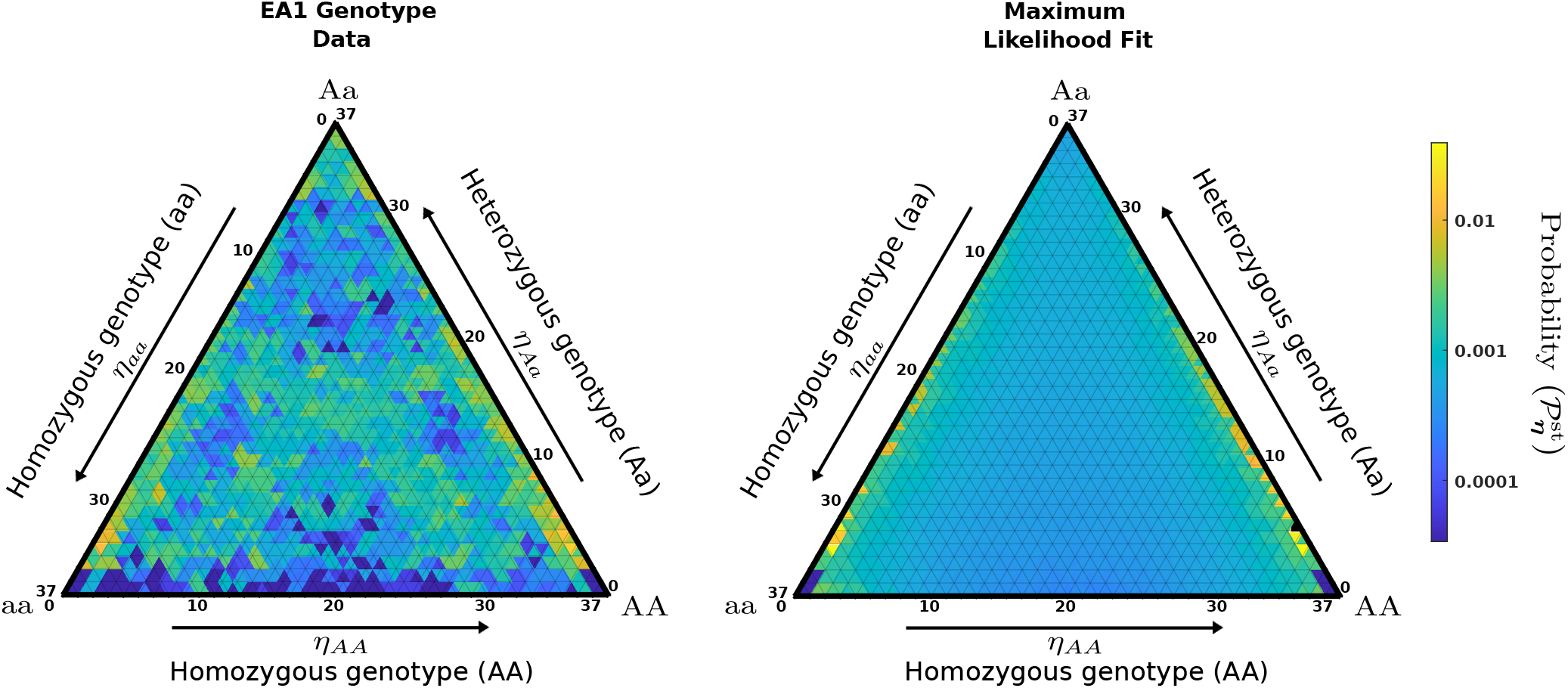
Comparison of *Leishmania* data for the EA1 population (left) with fitted model (right). Blue represents relatively low probability, whereas yellow represents relatively high probability of finding a locus at that position in the genotype space. Fitted parameters are given in Table 1. Ternary plots are produced using Lynch (2024). Analogous plots in cartesian coordinates can be found in Fig S7, along with corresponding likelihood surfaces.

We see that the model predicts that many of the populations appear highly inbred, with 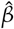close to one for EA2, ISC1 and ISC2.Meanwhile Brazil 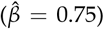 and the genetically diverse East African population 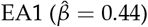 appear more outbred.

The number of estimated sexual events per generation, *N*_*f it*_*σ*_*f it*_ is generally similar across the populations, ranging from 1.8 for Brazil to 11.5 for EA2. However, the different magnitudes of the effective population sizes of the samples, *N*_*e*_, lead to a broader range in the estimates of the probability of sexual reproduction 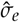 (see Eq. (12)). These estimates range from 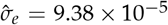 for EA1 (roughly equivalent to sexual reproduction once every 10^4^ generations) to 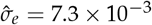 for ISC2 (roughly equivalent to sexual reproduction once every 140 generations).

The genome data we use in this analysis has some limitations. Some populations, such as EA1, EA2 and ISC2 have less than 50 samples. Given the challenges of obtaining parasites from VL endemic areas, and the complexity of parasite populations, this is unavoidable. Our analysis indicates that this limitation can lead to an underestimate for *σ* and overestimate for *β*, see Robustness checks on fitting. Additionally, some populations have very few segregating sites (e.g. ISC2 *M* < 1000), which can lead to a slight overestimate for *σ* and uncertainty in *β*, see Robustness checks on fitting. Although *M* > 1000 for EA1, EA2, ISC1 and Brazil, we have assumed that these sites are all independent (no linkage). Therefore information contained in the segregating sites *M* may be lower, which could lead to similar issues with estimates of *σ* and uncertainty in *β*. Finally, the scale of our error estimates in Table 1 assumes sites are all independent, so our estimates will be under-estimates.

## Discussion

Leishmaniasis is one of the most important neglected diseases, with between 1.5 to 2 million cases occurring each year (Den Boer *et al*. 2011). Drug treatment options are limited, toxic, and have high treatment failure rates, particularly for some clinical manifestations or geographic regions (Santos *et al*. 2023). No vaccine is available at this stage.

The ecology of *Leishmania*sis transmission is complicated by multiple sand fly vector species and mammal reservoirs (Mc-Dowell and Robichaud 2024). Population genetics has made an important contribution to understanding transmission and disease epidemiology (Schwabl *et al*. 2019; Cotton *et al*. 2020; Grace *et al*. 2022; Heeren *et al*. 2024). The frequency of sex, and the degree of inbreeding are particularly important since between-species hybrids appear to be frequent, and these can lead to new disease presentations and/or adaptation of parasite to new ecological settings (Van den Broeck *et al*. 2020; Lypaczewski and Matlashewski 2021). As meiosis only occurs within sand fly vectors, outbreeding and evidence for sex are indications of epidemiological circumstances where sand fly ‘residence time’ is important.

Outbreeding is also an indicator of transmission dynamics, since it indicates that parasites are not only shared within very local, inbred populations. Our analysis indicates that the *L. donovani* EA1 population from North Ethiopia and Sudan is very outbred (*β* = 0.44) while the *L. infantum* population in Brazil is relatively outbred (*β* = 0.77). This has implications for the spread of drug resistance alleles onto locally genetically adapted backgrounds.

From a theoretical standpoint, our approach to estimating the rate of sexual reproduction is similar to that of Stoeckel and Masson (2014a), in that it takes advantage of the departures from Hardy-Weinberg equilibrium (or modified equilibria that account for inbreeding) that are prevalent in populations with rare facultative sex. However in a departure from that work, we fitted data to the the full genotype distribution of the model rather than the distribution of Wright’s inbreeding coefficient, *F*_IS_. We were able to do this by exploiting the invariance of the genotype distribution for fixed *σN* (number of sexual events per generation) that has also been identified in other approaches (Hartfield 2016a; Hartfield *et al*. 2018; Ennos and Hu 2019).

The focus of our approach on a single locus has justification; in the rare sex regime, supervised machine learning has identified variance in *F*_IS_ as a key predictor of the rate of sexual reproduction (Stoeckel *et al*. 2021). However that study also identified the mean correlation coefficient of genetic distance between unordered alleles (a measure of linkage disequilibrium) as another key genetic index for predicting rates of rare sex. It would therefore be interesting to combine LD, as has been used successfully in other studies (Tsai *et al*. 2008b), with the approach we describe here to develop a pluralist method.

As with any modelling, there are of course elements that are missing from our model. For instance, we have not accounted for gene conversion which is prominent in fungal species (Halkett *et al*. 2005) and can have amplified effects when sex is rare (Hartfield 2016a; Hartfield *et al*. 2018). While gene conversion has a somewhat similar affect to inbreeding in our model in terms of reducing heterozygosity, it is interesting to note that the two processes would appear functionally distinct in the model (the gene conversion rate would be proportional to the number of heterozygotes, while the inbreeding rate is also proportional to homozygote frequencies, see Eq. (1)). It would therefore be interesting in future work to see if this could be leveraged to disentangle the two effects in *Leishmania* (Rogozin *et al*. 2020). Similarly we have also not accounted for the possibility of any self-incompatibility in *Leishmania*. While no mechanisms are currently known in *Leishmania*, the presence of such self-incompatible mating types has been used effectively to estimate *Nσ* in facultatively sexual fungi (Ennos and Hu 2019). Finally, haplotype selection (the loss of haplotypes resulting from unstable chromosome copies between generations) will also result in the loss of heterozygosity. This will cause populations to appear more inbred within chromosomes that have the most variation in their somy (Reis-Cunha *et al*. 2024b).

As with previous studies, we have assumed that the population is at drift–migration–mutation equilibrium (Stoeckel and Masson 2014a; Ennos and Hu 2019), as this is clearly the most analytically tractable scenario. However sampling a population that is out of equilibrium may also to lead to erroneous inferences (Reichel *et al*. 2016). In this respect, *Leishmania*, with its complex life-cycle, is an interesting case study. Since meiosis is restricted to occurring in insect vectors, populations are subject to temporally heterogeneous sexual reproduction large changes in cell numbers during development from from 10^2^ to 10^11^ cells, with frequent population bottlenecks (Kimblin *et al*. 2008; Verma *et al*. 2010; Serafim *et al*. 2018), which are not typically considered in the literature on estimating rates of sex (Hartfield 2016a), and indeed which we have not considered here. Incorporating such life-history traits would be challenging from a modelling perspective. However, the presence of “ecological designations” (i.e. where we expect organisms to be reproducing sexually) can be used in parallel with genetic studies to determine what reasonable estimates for rates of sexual reproduction might be Allen and Lynch (2012); Doerder *et al*. (1995).

While ecology can help constrain estimates of sexual reproduction rates, estimates of such rates can in turn inform epidemiology. For example, for the 20 *Leishmania* species that cause disease, differences in the frequency of sex are informative about the proportion of the life cycle that is within sand fly vectors (where meiosis occurs), compared to within mammalian hosts (where meiosis does not occur). Comparisons between the rate of sexual 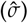 between disease foci in this way will generate information locations that greater sand fly vector parasite residence. Similarly, better estimates of inbreeding 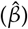 provide information about whether parasite populations are present in very isolated or regionally connected disease foci.

Despite these caveats the estimates of sexual reproduction we have obtained are consistent with other results for rare facultatively sexual organisms, including *Saccharomyces paradoxus* (*σ* ≈ 5 × 10^−4^), *Chlamydomonas reinhardtii* (*σ* ≈ 1.3 × 10^−3^), *Zymoseptoria tritici* (*Nσ* ⪆ 50), *Erisiphe necator* (*Nσ* ≈ 2 to 50), *Erisiphe necator* (*Nσ* ≈ 2 to > 50), *Rhyncosporium secalis* (*Nσ* ≈ 2.9 to > 50) and *Dothistroma septosporum* (*Nσ* ≈ 4.2 to 22.2) (Tsai *et al*. 2008a; Hasan and Ness 2018; Ennos and Hu 2019).

In this paper we have provided the first estimate for the rate of sexual reproduction in the *Leishmania*. With the five *Leishmania donovani* species complex populations we examine, we observe considerable outbreeding in two populations (East Africa and Brazil). All other populations appear much more inbred, but our analysis shows that limitations in sample sizes and/or segregating sites can result in inaccurate estimates of 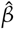.Frequencies of sex of the order of one sexual reproduction per 10^3^ - 10^4^ mitotic generations appear reasonable given our understanding of the number of cells present in mammalian host compared to sand flies.

It has been long been argued that methods to infer rates of sexual reproduction using population genetics models required further development (Schurko *et al*. 2009; Birky Jr 2010). Such methods are crucial in facultatively sexual pathogens (de Meeûs *et al*. 2007). In the particular case of *Leishmania*, not only can this quantity help to understand *Leishmania* evolution, including the spread of drug resistance and the eco-epidemiological dynamics of parasite populations in different contexts.

## Data availability

Genotype data from *Leishmania donovani* populations was derived from Grace *et al*. (2021) and is available in VCF format at https://figshare.com/projects/Balancing_selection_in_Leishmania_donovani/94292. Code to generate parameter estimates in Table 1 is available on request.

## Acknowledgments

This project utilised the Viking Research computing cluster at the University of York. We acknowledge the support of The Genomics Laboratory in the Technology Facility, Department of Biology, University of York.

