## Supplemental Information for "Estimating the rate of sexual reproduction and the inbreeding rate in *Leishmania*"

#### Contents

**S1 Table of parameters** **S3**

**S2 Mathematical Model** **S4**

S2.1 Transition Rates and stationary distribution of the master equation . . . . . . . . . . . . S4

S2.2 Derivation of the Fokker-Planck equation . . . . . . . . . . . . . . . . . . . . . . . . . . S6

S2.3 Deterministic Dynamics . . . . . . . . . . . . . . . . . . . . . . . . . . . . . . . . . . . S7

S2.4 The compound parameter $N\sigma$ in the rare-sex diffusive limit . . . . . . . . . . . . . . . . . S9

**S3 Empirical Data** **S10**

S3.1 Minor allele frequency filtering . . . . . . . . . . . . . . . . . . . . . . . . . . . . . . . S10

S3.2 Estimation of the effective population size $N_e$ . . . . . . . . . . . . . . . . . . . . . . . S10

**S4 Fitting simulated data and robustness checking** **S11**

S4.1 Example: Fitting the stationary distribution $P^{\text{st}}_{\boldsymbol{n}}$ to simulated data . . . . . . . . . . . . . S11

S4.2 Fitting the sampled stationary distribution $\mathcal{P}^{\text{st}}_{\boldsymbol{\eta}}$ to simulated data using $N\sigma$ Theory . . . S13

S4.2.1 Simulated Data . . . . . . . . . . . . . . . . . . . . . . . . . . . . . . . . . . . . . S16

S4.2.2 Varying the model fitting population size, $N_{fit}$ . . . . . . . . . . . . . . . . . . . . S17

S4.2.3 Varying the number of observations used in model fitting, $M$ . . . . . . . . . . . . S19

S4.2.4 Varying the number of individuals sampled from the wider population, $N_s$ . . . . S21

[*]Department of Mathematics, University of York

[†]School of Biological Sciences, University of Edinburgh

[‡]Department of Physics and Astronomy, University of Exeter

[§]Department of Biology and York Biomedical Research Institute, University of York

[2]Equal contribution

**S5 Importance of co-fitting inbreeding rate and rate of sexual reproduction** **S23**

S5.1 Estimation of Wright's inbreeding coefficient using $\beta$ . . . . . . . . . . . . . . . . . . . . . S23

S5.2 Estimation of Wright's Inbreeding coefficient using expected and observed heterozygosity S23

S5.3 Estimation of the rate of facultative sexual reproduction in the effective population $\sigma_e$ . . S24

**S6 Results of model fitting** **S25**

S6.1 EA1 . . . . . . . . . . . . . . . . . . . . . . . . . . . . . . . . . . . . . . . . . . . . . S26

S6.2 EA2 . . . . . . . . . . . . . . . . . . . . . . . . . . . . . . . . . . . . . . . . . . . . . S27

S6.3 ISC1 . . . . . . . . . . . . . . . . . . . . . . . . . . . . . . . . . . . . . . . . . . . . . S28

S6.4 ISC2 . . . . . . . . . . . . . . . . . . . . . . . . . . . . . . . . . . . . . . . . . . . . . S29

S6.5 Brazil . . . . . . . . . . . . . . . . . . . . . . . . . . . . . . . . . . . . . . . . . . . . S30

**S7 de Finetti Diagrams** **S31**

#### S1 Table of parameters

| Symbol | Description |
|---|---|
| $N$ | Population size |
| $\sigma$ | Rate of facultative sexual reproduction |
| $\beta$ | Rate of inbreeding |
| $T$ | Total simulation time |
| $n_{AA}$ | Number of individuals with homozygous $AA$ genotype |
| $n_{Aa}$ | Number of individuals with heterozygous $Aa$ genotype |
| $n_{aa}$ | Number of individuals with homozygous $aa$ genotype |
| $x_{AA}$ | Frequency of individuals with homozygous $AA$ genotype |
| $x_{Aa}$ | Frequency of individuals with heterozygous $Aa$ genotype |
| $x_{aa}$ | Frequency of individuals with homozygous $aa$ genotype |
| $N_{samp}$ | Sample population size |
| $N_e$ | Effective population size |
| $\hat{N}_e$ | Estimate of the effective population size |
| $N_{\text{fit}}$ | Population size used as part of the fitting algorithm |
| $\hat{\sigma}$ | Maximum likelihood estimate of the rate of facultative sexual reproduction |
| $\hat{\sigma}_{fit}$ | Maximum likelihood estimate of the scaled rate of facultative sexual reproduction |
| $\hat{\sigma}_e$ | Maximum likelihood estimate of the effective rate of facultative sexual reproduction |
| $\hat{\beta}$ | Maximum likelihood estimate of the rate of inbreeding |
| $F_{IS}$ | Wright's Inbreeding coefficient |
| $F_{IS}(\beta)$ | Estimate of Wright's Inbreeding coefficient using inbreeding rate, see Eq. (S57). |
| $\beta_{\text{het}}(F_{IS})$ | Estimate of inbreeding rate from Wright's inbreeding coefficient, see Eq. (S58). |

Table S1: Table of parameters and variables used in the 2D diploid Moran model with inbreeding and facultative sexual reproduction and their descriptions.

### S2 Mathematical Model

#### S2.1 Transition Rates and stationary distribution of the master equation

The six general transition rates are

$$\begin{aligned}T_1(\boldsymbol{n}) = T[(n_{AA}+1, n_{Aa})|(n_{AA}, n_{Aa})] = \Bigg[\sigma\Bigg((1-\beta)\left[\frac{n_{AA}^2}{N^2} + \frac{n_{AA}n_{Aa}}{N^2} + \frac{1}{4}\frac{n_{Aa}^2}{N^2}\right] + \beta\left[\frac{n_{AA}}{N} + \frac{1}{4}\frac{n_{Aa}}{N}\right]\Bigg) \\ + (1-\sigma)\left(\frac{n_{AA}}{N}\right)\Bigg]\frac{(N-n_{AA}-n_{Aa})}{N}\end{aligned} \tag{S1}$$

$$\begin{aligned}T_2(\boldsymbol{n}) = T[(n_{AA}-1, n_{Aa})|(n_{AA}, n_{Aa})] = \Bigg[\sigma\Bigg((1-\beta)\left[\frac{1}{4}\frac{(n_{Aa})^2}{N^2} + \frac{n_{Aa}(N-n_{AA}-n_{Aa})}{N^2} + \frac{(N-n_{AA}-n_{Aa})^2}{N^2}\right] \\ + \beta\left[\frac{(N-n_{AA}-n_{Aa})}{N} + \frac{1}{4}\frac{n_{Aa}}{N}\right]\Bigg) \\ + (1-\sigma)\left(\frac{(N-n_{AA}-n_{Aa})}{N}\right)\Bigg]\frac{n_{AA}}{N}\end{aligned} \tag{S2}$$

$$\begin{aligned}T_3(\boldsymbol{n}) = T[(n_{AA}, n_{Aa}+1)|(n_{AA}, n_{Aa})] = \Bigg[\sigma\Bigg((1-\beta)\Bigg[\frac{n_{AA}n_{Aa}}{N^2} + \frac{2n_{AA}(N-n_{AA}-n_{Aa})}{N^2} + \frac{1}{2}\frac{(n_{Aa})^2}{N^2} \\ + \frac{n_{Aa}(N-n_{AA}-n_{Aa})}{N^2}\Bigg] + \beta\left[\frac{1}{2}\frac{n_{Aa}}{N}\right]\Bigg) \\ + (1-\sigma)\left(\frac{n_{Aa}}{N}\right)\Bigg]\frac{(N-n_{AA}-n_{Aa})}{N}\end{aligned} \tag{S3}$$

$$\begin{aligned}T_4(\boldsymbol{n}) = T[(n_{AA}, n_{Aa}-1)|(n_{AA}, n_{Aa})] = \Bigg[\sigma\Bigg((1-\beta)\left[\frac{1}{4}\frac{(n_{Aa})^2}{N^2} + \frac{n_{Aa}(N-n_{AA}-n_{Aa})}{N^2} + \frac{(N-n_{AA}-n_{Aa})^2}{N^2}\right] \\ + \beta\left[\frac{(N-n_{AA}-n_{Aa})}{N} + \frac{1}{4}\frac{n_{Aa}}{N}\right]\Bigg) \\ + (1-\sigma)\left(\frac{(N-n_{AA}-n_{Aa})}{N}\right)\Bigg]\frac{n_{Aa}}{N}\end{aligned} \tag{S4}$$

$$\begin{aligned}T_5(\boldsymbol{n}) = T[(n_{AA}+1, n_{Aa}-1)|(n_{AA}, n_{Aa})] = \Bigg[\sigma\Bigg((1-\beta)\left[\frac{(n_{AA})^2}{N^2} + \frac{n_{AA}n_{Aa}}{N^2} + \frac{1}{4}\frac{(n_{Aa})^2}{N^2}\right] \\ + \beta\left[\frac{n_{AA}}{N} + \frac{1}{4}\frac{n_{Aa}}{N}\right]\Bigg) + (1-\sigma)\left(\frac{n_{AA}}{N}\right)\Bigg]\frac{n_{Aa}}{N}\end{aligned} \tag{S5}$$

$$\begin{aligned}T_6(\boldsymbol{n}) = T[(n_{AA}-1, n_{Aa}+1)|(n_{AA}, n_{Aa})] = \Bigg[\sigma\Bigg((1-\beta)\Bigg[\frac{n_{AA}n_{Aa}}{N^2} + \frac{2n_{AA}(N-n_{AA}-n_{Aa})}{N^2} + \frac{1}{2}\frac{(n_{Aa})^2}{N^2} \\ + \frac{n_{Aa}(N-n_{AA}-n_{Aa})}{N^2}\Bigg] + \beta\left[\frac{1}{2}\frac{n_{Aa}}{N}\right]\Bigg) \\ + (1-\sigma)\left(\frac{n_{Aa}}{N}\right)\Bigg]\frac{n_{AA}}{N}\end{aligned} \tag{S6}$$

Together with the master equation, Eq. (2) the main text, the transition rates determine the stochastic dynamics of the model. Since Eq. (2) is linear, it can be written more compactly in matrix notation;

$$\frac{d\boldsymbol{P}}{dt} = \boldsymbol{M}\boldsymbol{P}\,. \tag{S7}$$

where $P$ is a vector of length $(N+1)(N+2)/2$ with elements corresponding to each of the $(N+1)(N+2)/2$ possible states $(n_{AA}, n_{Aa})$. The matrix $M$ meanwhile is a square $[(N+1)(N+2)/2] \times [(N+1)(N+2)/2]$ matrix. From the form of Eq. (S7), we can show that the sum of each column must sum to zero [1]

$$\sum_{i \neq j} M_{ij} = -M_{ii}\,, \tag{S8}$$

which follows from the fact that probability is conserved (transitions out of some state $i^{\text{th}}$ state must end up in some other state $j^{\text{th}}$ state). This structure ensures that $M$ has at least one zero eigenvalue.

The stationary probability distribution $\boldsymbol{P}^{\text{st}}$ can be obtained by taking the limit of $t \to \infty$ in Eq. (S7) (setting the derivative on the left-hand side of Eq. (S7) to zero). Finding the stationary distribution is then equivalent to an eigenproblem

$$\boldsymbol{M}\boldsymbol{P}^{st} = \boldsymbol{0} \tag{S9}$$

in which solutions $\boldsymbol{P}^{st}$ are eigenvectors corresponding to zero eigenvalue. If the general transitions rates in Eqs. (S1-S6) are used for all $n_{AA} \in [0, N]$ and $n_{Aa} \in [0, N]$ then there are two such solutions for $\boldsymbol{P}^{st}$ (i.e. two eigenvalues of $M$) corresponding to the two absorbing states $(n_{AA}, n_{Aa}) = (0, 0)$ and $(n_{AA}, n_{Aa}) = (N, 0)$ (see red states in Figure 1 in the main text).

In the Mathematical Modeling section of the main text, we describe why we are not interested in these absorbing states, which correspond to the extinction of allele $a$ and allele $A$ respectively. We therefore alter the transitions in Eqs. (S1-S6), such that they apply to all initial states $\boldsymbol{n}$ except $\boldsymbol{n} = (1, 0)$ and $\boldsymbol{n} = (N-1, 0)$ (see modified blue transitions in Figure 1 in the main text). This is equivalent to setting up reflecting boundaries at the absorbing states. For transitions out of state $\boldsymbol{n} = (1, 0)$, we now have

$$\begin{aligned}
T_1(\boldsymbol{n}) = T(2,0|1,0) &= \frac{1}{N^3}(N-1)\left[N - \sigma(1-\beta)(N-1)\right], && \text{(S10)}\\
T_2(\boldsymbol{n}) = T(0,0|1,0) &= 0, && \text{(S11)}\\
T_3(\boldsymbol{n}) = T(1,1|1,0) &= \frac{1}{N^3}2(N-1)^2\sigma(1-\beta), && \text{(S12)}\\
T_4(\boldsymbol{n}) = T(1,-1|1,0) &= 0, && \text{(S13)}\\
T_5(\boldsymbol{n}) = T(2,-1|1,0) &= 0, && \text{(S14)}\\
T_6(\boldsymbol{n}) = T(0,1|1,0) &= \frac{1}{N^3}\left[2(N-1)\sigma(1-\beta) + (N-1)(N-\sigma(1-\beta))\right], && \text{(S15)}
\end{aligned}$$

where altered terms in the transitions (i.e. that set $T_2(1, 0)$ to zero and increase $T_6(1, 0)$ by the corresponding amount) have been given in red (set to zero) and blue (terms added to alternative transition). Meanwhile for $\boldsymbol{n} = (N-1, 0)$ we have

$$\begin{aligned}
T_1(\boldsymbol{n}) = T(N,0|N-1,0) &= 0, && \text{(S16)}\\
T_2(\boldsymbol{n}) = T(N-2,0|N-1,0) &= \frac{1}{N^3}(N-1)\left[N - \sigma(1-\beta)(N-1)\right], && \text{(S17)}\\
T_3(\boldsymbol{n}) = T(N-1,1|N-1,0) &= \frac{1}{N^3}\left[2(N-1)\sigma(1-\beta) + (N-1)(N-\sigma(1-\beta))\right], && \text{(S18)}\\
T_4(\boldsymbol{n}) = T(N-1,-1|N-1,0) &= 0, && \text{(S19)}\\
T_5(\boldsymbol{n}) = T(N-1,1|N-1,0) &= 0, && \text{(S20)}\\
T_6(\boldsymbol{n}) = T(N-2,1|N-1,0) &= \frac{1}{N^3}2(N-1)^2\sigma(1-\beta), && \text{(S21)}
\end{aligned}$$

where altered terms in the transitions (i.e. that set $T_1(N-1,0)$ to zero and increase $T_3(N-1,0)$ by the corresponding amount) have been given in red (set to zero) and blue (terms added to alternative transition).

Once the matrix $M$ is amended to include Eqs. (S10-S21), the matrix features just one zero eigenvalue. Therefore there is just one stationary state, given by the eigenvector of $M$ associated with this zero eigenvalue.

#### S2.2 Derivation of the Fokker-Planck equation

Assuming $N$ is large, we can use the diffusion approximation [2] to obtain an approximation for the probabilistic dynamics in the approximately continuous variable $\boldsymbol{x} = \boldsymbol{n}/N$ (i.e. we can conduct a change of variables $\boldsymbol{n} = \boldsymbol{x}N$, followed by a Taylor expansion of Eq. (2) in the small parameter $1/N$, also known as the Kramers-Moyal Expansion [3]). We obtain the following Fokker-Planck equation (FPE):

$$\frac{\partial p(\boldsymbol{x},t)}{\partial t} = -\frac{1}{N}\sum_{i=1}^{m}\frac{\partial}{\partial x_i}\underbrace{[A_i(\boldsymbol{x})p(\boldsymbol{x},t)]}_{\text{Advection term}} + \frac{1}{2N^2}\sum_{i,j=1}^{m}\frac{\partial^2}{\partial x_i \partial x_j}\overbrace{[B_{ij}(\boldsymbol{x})p(\boldsymbol{x},t)]}^{\text{Diffusion term}} \tag{S22}$$

where $\boldsymbol{A}(\boldsymbol{x})$ is the advection vector and $\boldsymbol{B}(\boldsymbol{x})$ is the diffusion, or noise, matrix. In general the advection vector and diffusion matrix are given by [4]

$$A_i(\boldsymbol{x}) = \lim_{N\to\infty}\sum_{k=1}^{u}\nu_{ik}f_k(\boldsymbol{x}) \tag{S23}$$

and

$$B_{ij}(\boldsymbol{x}) = \lim_{N\to\infty}\sum_{k=1}^{u}\nu_{ik}\nu_{jk}f_k(\boldsymbol{x})\,, \tag{S24}$$

where $\nu$ is the stoichiometry matrix and $f_i(\boldsymbol{x})$ are continuous approximations of the probability transition rates, such that $f_i(\boldsymbol{x}) = T_i(\boldsymbol{x}N)|_{N\to\infty}$. For the problem at hand, we have

$$\nu = \begin{pmatrix} 1 & -1 & 0 & 0 & 1 & -1 \\ 0 & 0 & 1 & -1 & -1 & 1 \end{pmatrix}. \tag{S25}$$

Therefore the elements of the advection term are given by

$$A_{x_{AA}}(\boldsymbol{x}) = -\sigma\big[(\beta-1)(x_{AA})^2 + (\beta-1)(x_{Aa}-1)x_{AA} + \frac{1}{4}x_{Aa}\left((\beta-1)x_{Aa}-\beta\right)\big]\,, \tag{S26}$$

$$A_{x_{Aa}}(\boldsymbol{x}) = 2\,\sigma\big[(\beta-1)(x_{AA})^2 + (\beta-1)(x_{Aa}-1)x_{AA} + \frac{1}{4}x_{Aa}\left((\beta-1)x_{Aa}-\beta\right)\big]\,, \tag{S27}$$

and the elements of the diffusion matrix by

$$
\begin{aligned}
B_{x_{AA}x_{AA}}(\boldsymbol{x}) &= 2\sigma(\beta-1)(x_{AA}(t))^3 + \frac{(8\sigma(\beta-1)x_{Aa}(t) - 8 + (-12\beta+12)\sigma)(x_{AA}(t))^2}{4} \\
&+ \frac{(2\sigma(\beta-1)(x_{Aa}(t))^2 + (-6\beta+4)\sigma x_{Aa}(t) + 8 + \sigma(4\beta-4))x_{AA}(t)}{4} \\
&- \frac{((\beta-1)x_{Aa}(t)-\beta)x_{Aa}(t)\sigma}{4}\,, \qquad \text{(S28)} \\
B_{x_{AA}x_{Aa}}(\boldsymbol{x}) = B_{x_{Aa}x_{AA}}(\boldsymbol{x}) &= -2\sigma(\beta-1)(x_{AA}(t))^3 - \sigma(x_{Aa}(t)-2)(\beta-1)(x_{AA}(t))^2 \\
&+ \frac{x_{Aa}(t)(\sigma(\beta-1)x_{Aa}(t) - 4 + (-\beta+2)\sigma)x_{AA}(t)}{2} \\
&+ \frac{((\beta-1)x_{Aa}(t)-\beta)(x_{Aa}(t))^2\sigma}{4}\,, \qquad \text{(S29)} \\
B_{x_{Aa}x_{Aa}}(\boldsymbol{x}) &= -\sigma(\beta-1)(x_{Aa}(t))^3 \\
&+ \frac{(-8\sigma(\beta-1)x_{AA}(t) - 4 + (3\beta-1)\sigma)(x_{Aa}(t))^2}{2} \\
&+ \frac{(-8\sigma(\beta-1)(x_{AA}(t))^2 + 12\sigma(\beta-1)x_{AA}(t) - \sigma\beta + 4)x_{Aa}(t)}{2} \\
&+ 2\sigma x_{AA}(t)(x_{AA}(t)-1)(\beta-1)\,. \qquad \text{(S30)}
\end{aligned}
$$

#### S2.3 Deterministic Dynamics

In the infinite population size limit, the dynamics of the geneotype frequencies $x_{AA} = n_{AA}/N$ and $x_{Aa} = n_{Aa}/N$ are given by [5]

$$\frac{\mathrm{d}\boldsymbol{x}}{\mathrm{d}t} = \boldsymbol{A}(\boldsymbol{x}) \qquad \text{(S31)}$$

or

$$
\begin{aligned}
\frac{\mathrm{d}x_{AA}}{\mathrm{d}\tau} &= -\,\sigma\big[(\beta-1)(x_{AA})^2 + (\beta-1)(x_{Aa}-1)x_{AA} + \frac{1}{4}x_{Aa}\left((\beta-1)x_{Aa}-\beta\right)\big] \qquad \text{(S32)} \\
\frac{\mathrm{d}x_{Aa}}{\mathrm{d}\tau} &= 2\,\sigma\big[(\beta-1)(x_{AA})^2 + (\beta-1)(x_{Aa}-1)x_{AA} + \frac{1}{4}x_{Aa}\left((\beta-1)x_{Aa}-\beta\right)\big]. \qquad \text{(S33)}
\end{aligned}
$$

Eqs. (S32-S33) feature a line of fixed points (along which $\mathrm{d}x_{AA}/\mathrm{d}\tau = 0$ and $\mathrm{d}x_{Aa}/\mathrm{d}\tau = 0$) to which the system relaxes at long times (see red dashed line in Main Text Figure 2, left panels). This line of fixed points is termed the centre manifold, and can be expressed as

$$x_{Aa} = \frac{-\gamma + 4x_{AA} + \sqrt{\gamma^2 + 8\gamma x_{AA} + 16x_{AA}}}{2}\,, \qquad \text{(S34)}$$

with

$$\gamma = \frac{\beta}{1-\beta}\,. \qquad \text{(S35)}$$

For no inbreeding ($\beta = 0$) Eq. (S44) reduces to

$$x_{Aa} = 2(\sqrt{x_{AA}} - x_{AA})\,. \qquad \text{(S36)}$$

This is simply the Hardy-Weinberg equilibrium. One can see this by noting that for the standard Hardy-Weinberg equation with $q_A$ for the frequency of allele $A$ and $q_a$ for the frequency of allele $a$, we have

$$q_A^2 + 2q_A q_a + q_a^2 = 1\,, \tag{S37}$$

with homozygous frequencies $x_{AA} = q_A^2$ and $x_{aa} = q_a^2 = (1 - x_{AA} - x_{Aa})$, and heterozygous frequencies $x_{Aa} = 2q_a q_A$. Solving

$$x_{AA} = q_A^2\big|_{q_A = \frac{2x_{AA}+x_{Aa}}{2}} \tag{S38}$$

for $x_{AA}$, we recover Eq. (S36), and similarly for $x_{Aa} = 2q_a q_A$.

In the case where the inbreeding is present ($\beta > 0$), we must instead compare Eq. (S44) to the Hardy-Weinberg equilibrium modified to account for inbreeding. We have [6]:

$$(1 - F_{\mathrm{IS}})q_A^2 + F_{\mathrm{IS}}q_A + (1 - F_{\mathrm{IS}})2q_A q_a + (1 - F_{\mathrm{IS}})q_a^2 + F_{\mathrm{IS}}q_a = 1\,, \tag{S39}$$

where $F_{\mathrm{IS}}$ is Wright's inbreeding coefficient. Here the frequency of homozygotes is now given by $x_{AA} = (1 - F_{\mathrm{IS}})q_A^2 + F_{\mathrm{IS}}q_A$ and $x_{aa} = (1 - F_{\mathrm{IS}})q_a^2 + F_{\mathrm{IS}}q_a$, and the frequency of heterozygotes by $x_{Aa} = (1 - F_{\mathrm{IS}})2q_a q_A$ (i.e. heterzygote frequency is reduced under inbreeding). Now solving

$$x_{AA} = (1 - F_{\mathrm{IS}})q_A^2 + F_{\mathrm{IS}}q_A\big|_{q_A = \frac{2x_{AA}+x_{Aa}}{2}} \tag{S40}$$

for $x_{AA}$, we recover Eq. (S44) so long as

$$F_{\mathrm{IS}} = \frac{\beta}{2 - \beta}\,, \tag{S41}$$

as shown by [7]. This now gives a way to relate the inbreeding probability $\beta$ to Wright's inbreeding coefficient.

Alternatively one can note that Wright's inbreeding coefficient is defined as

$$F_{IS} = \frac{\langle H_{exp}\rangle - \langle H_{obs}\rangle}{\langle H_{exp}\rangle}\,, \tag{S42}$$

where $\langle H_{exp}\rangle$ is the expected heterozygosity under no inbreeding (i.e. at Hardy-Weinberg equilibrium) and $\langle H_{obs}\rangle$ is the observed heterozygosity. In order to couch this in terms of our model, we begin by writing $x_{Aa}$ as a function of $q_A$ on the center-manifold given by . Solving $q_A = (2x_{AA} + x_{Aa})/2$ for $x_A A$, we obtain $x_{AA} = (2q_A - x_{Aa})/2$. Now solving

$$x_{Aa} = \left.\frac{-\gamma + 4x_{AA} + \sqrt{\gamma^2 + 8\gamma x_{AA} + 16x_{AA}}}{2}\right|_{x_{AA}=(2q_A - x_{Aa})/2}\,, \tag{S43}$$

for $x_{Aa}$, we obtain

$$x^*_{Aa} = 4q_A(1-q_a)\frac{(1-\beta)}{(2-\beta)}\,. \tag{S44}$$

For the model, Eq. (S59) can then be expressed

$$\begin{aligned} F_{IS} &= 1 - \frac{x^*_{Aa}}{x^*_{Aa}|_{\beta=0}}\,, \\ &= 1 - 2\frac{(1-\beta)}{(2-\beta)}\,, \\ &= \frac{\beta}{2-\beta}\,. \end{aligned}$$

#### S2.4 The compound parameter $N\sigma$ in the rare-sex diffusive limit

We begin by re-writing Eqs. (S26-S30) to capture their dependence on $\sigma$ at different orders. For the advection vector we find

$$\boldsymbol{A}(\boldsymbol{x}) = \sigma \boldsymbol{A}^{(\sigma)}(\boldsymbol{x})\,, \tag{S45}$$

while for the diffusion matrix we have

$$\boldsymbol{B}(\boldsymbol{x}) = \boldsymbol{B}^{(1)}(\boldsymbol{x}) + \sigma \boldsymbol{B}^{(\sigma)}(\boldsymbol{x})\,. \tag{S46}$$

The Taylor expansion in $1/N$ (or diffusion approximation) used to obtain the FPE in Eq. (S22) yields

$$\frac{\partial p(\boldsymbol{x},t)}{\partial t} = -\frac{\sigma}{N}\sum_{i=1}^{2}\frac{\partial}{\partial x_i}\left[A_i^{(\sigma)}(\boldsymbol{x})p(\boldsymbol{x},t)\right] + \frac{1}{2N^2}\sum_{i,j=1}^{2}\frac{\partial^2}{\partial x_i \partial x_j}\left[\left(B_{ij}^{(1)}(\boldsymbol{x}) + \sigma B_{ij}^{(\sigma)}(\boldsymbol{x})\right)p(\boldsymbol{x},t)\right] + \mathcal{O}\left(\frac{1}{N^3}\right) \tag{S47}$$

where we note that order $N^{-3}$ terms have been truncated in Eq. (S22). Should the rate of sexual reproduction be small, such that $\sigma \approx \mathcal{O}(N^{-1})$ (i.e. of the order of a few sexual reproduction events per $N$ reproductive events), the equation simplifies to

$$\frac{\partial p(\boldsymbol{x},t)}{\partial t} = -\frac{\sigma}{N}\sum_{i=1}^{2}\frac{\partial}{\partial x_i}\left[A_i^{(\sigma)}(\boldsymbol{x})p(\boldsymbol{x},t)\right] + \frac{1}{2N^2}\sum_{i,j=1}^{2}\frac{\partial^2}{\partial x_i \partial x_j}\left[\left(B_{ij}^{(1)}(\boldsymbol{x})\right)p(\boldsymbol{x},t)\right] + \mathcal{O}\left(\frac{1}{N^3}\right)\,. \tag{S48}$$

Truncating the higher order terms in this equation, and rearranging for the stationary distribution $p^{st}(\boldsymbol{x})$ (with $\partial p^{st}(\boldsymbol{x})/\partial t = 0$), we find $p^{st}(\boldsymbol{x})$ is the solution to

$$\sum_{i,j=1}^{2}\frac{\partial^2}{\partial x_i \partial x_j}\left[B_{ij}^{(1)}(\boldsymbol{x})p^{st}(\boldsymbol{x})\right] = 2\sigma N\sum_{i=1}^{2}\frac{\partial}{\partial x_i}\left[A_i^{(\sigma)}(\boldsymbol{x})p^{st}(\boldsymbol{x})\right]\,, \tag{S49}$$

as given in Eq (5) of the main text.

### S3 Empirical Data

For full details of the data acquisition for the *Leishmania* populations used in this article see [8].

#### S3.1 Minor allele frequency filtering

As described in [8], the population data are filtered to exclude all loci where the frequency of the minor allele (MAF) was less than 0.05. Mathematically, this is equivalent to sampling from the region of state space

$$0.05 < \frac{2n_{AA} + n_{Aa}}{2N} < 0.95 \tag{S50}$$

in our model. We further excluded data arrising from chromosome 31 due to its consistent tetraploidy in *Leishmania*, which is inconsistent with our diploid model.

#### S3.2 Estimation of the effective population size $N_e$

Part of the calculation in the section Parameter Fitting of the main text (for obtaining an estimate of the rate of facultative sexual reproduction in the population, $\hat{\sigma}_e$) involves an estimate of the effective population size, $\hat{N}_e$ (see Eq. (12) and Table 1). This can be calculated using the following relationship between the nucleotide diversity ($\theta$) and the genome wide mutation rate in a diploid population ($\mu$) [9]:

$$\theta = 4N_e\mu\,. \tag{S51}$$

Using an estimate of nucleotide diversity ($\hat{\pi}$), $N_e$ can be estimated using the following formula:

$$\hat{N}_e = \frac{\hat{\pi}}{4\hat{\mu}}. \tag{S52}$$

Substituting values of $\hat{\pi}$ estimated from these populations of Leishmania (see Table in [8]) along with an estimate of the genome-wide mutation rate $\hat{\mu} = 1.99 \times 10^{-9}$ from [10], into Eq (S52), returns the estimates $\hat{N}_e$ seen in Table S2 and Table 1 in the main text.

| **Data set** | $\hat{\pi}$ ($\times 10^{-6}$) | $\hat{N}_e$ |
|---|---|---|
| EA1 | 424 | 53266 |
| EA2 | 87 | 10930 |
| ISC1 | 84 | 10553 |
| ISC2 | 6.3 | 791 |
| Brazil | 28 | 3518 |

Table S2: Estimates of the effective population size $\hat{N}_e$ of several populations of Leishmania obtained using Eq (S52). The estimates of nucleotide diversity $\hat{\pi}$ have been taken from Table in [8] and the mutation rate $\hat{\mu} = 1.99 \times 10^{-9}$ from [10].

### S4 Fitting simulated data and robustness checking

In this section we conduct a robustness check for our approach using simulated data. We begin in Section S4.1 fitting the simulation data using the full genotype distribution $P_{\boldsymbol{n}}^{\mathrm{st}}$ (see Eq. (3) of the main text) as a proof of principle. We follow this in Section S4.2 by showing that the sampled stationary distribution, $\mathcal{P}_{\boldsymbol{\eta}}^{\mathrm{st}}(N_s, N, \sigma, \beta)$ (defined in Eq. (7) of the main text), is unchanged if $N\sigma$ is held constant. In Section S4.2.2 we illustrate how this property can be exploited to speed up the fitting process. In Section S4.2.3 we explore the effect of reducing the number of "observations" of the state space (equivalent to reducing the number of segregating alleles observed in the population). Finally in Section S4.2.4 we explore the effect of reducing the number of individuals sampled from the population.

#### S4.1 Example: Fitting the stationary distribution $P_{\boldsymbol{n}}^{\mathrm{st}}$ to simulated data

To verify that the model fitting algorithm could correctly distinguish data across a range of parameter regimes, simulated data was generated according to a combination of different parameter values. Four independent simulated data sets were generated using a population $N_{sim} = 100$ and the following combination of parameters: $\sigma = 0.1$, and $\beta = 0.0$ (10% facultative sex and no inbreeding); $\sigma = 0.1$, and $\beta = 0.9$ (10% facultative sex and high inbreeding); $\sigma = 1.0$, and $\beta = 0.0$ (obligate sex and no inbreeding); $\sigma = 1.0$, and $\beta = 0.9$ (obligate sex and high inbreeding). All initial conditions were $(n_{AA}, n_{Aa}) = (33, 34)$. The total run time of each simulation was approximately $T = 1.3 \times 10^7$ generations (approximately $1.3 \times 10^9$ birth-death events), with no burn in time. Data was sampled (on average) every 100 generations (i.e. every $100 \times N_{\mathrm{sim}}$ timesteps) and filtered according to Eq. (S50), resulting in approximately $1.3 \times 10^9$ observations in total for each data set. This data can be found in [11]. Figure S1 illustrates the capability of the fitting algorithm to distinguish between simulated data sets generated using different parameter regimes. Model parameters were varied such that $0 \leq \beta \leq 0.9$ and $0.1 \leq \sigma \leq 1.0$, at intervals of 0.1, producing a grid of parameters which were used to construct models that were fitted to each data set. The log-likelihood was calculated for each model with respect to each data set, returning the maximum likelihood parameter estimates. As seen in Figure S1, for each parameter regime, the model fitting returns the exact combination of $\sigma$ and $\beta$ that were used to generate the data, for each data set. Therefore we conclude that in principle fitting data to $P_{\boldsymbol{n}}^{\mathrm{st}}$ can be used to distinguish between different parameter regimes and estimate parameters in simulated data. However real data typically features samples from populations with large sizes that are too great to allow for the efficient numerical solution of $P_{\boldsymbol{n}}^{\mathrm{st}}$. We therefore introduce a modified approach in the next subsection.

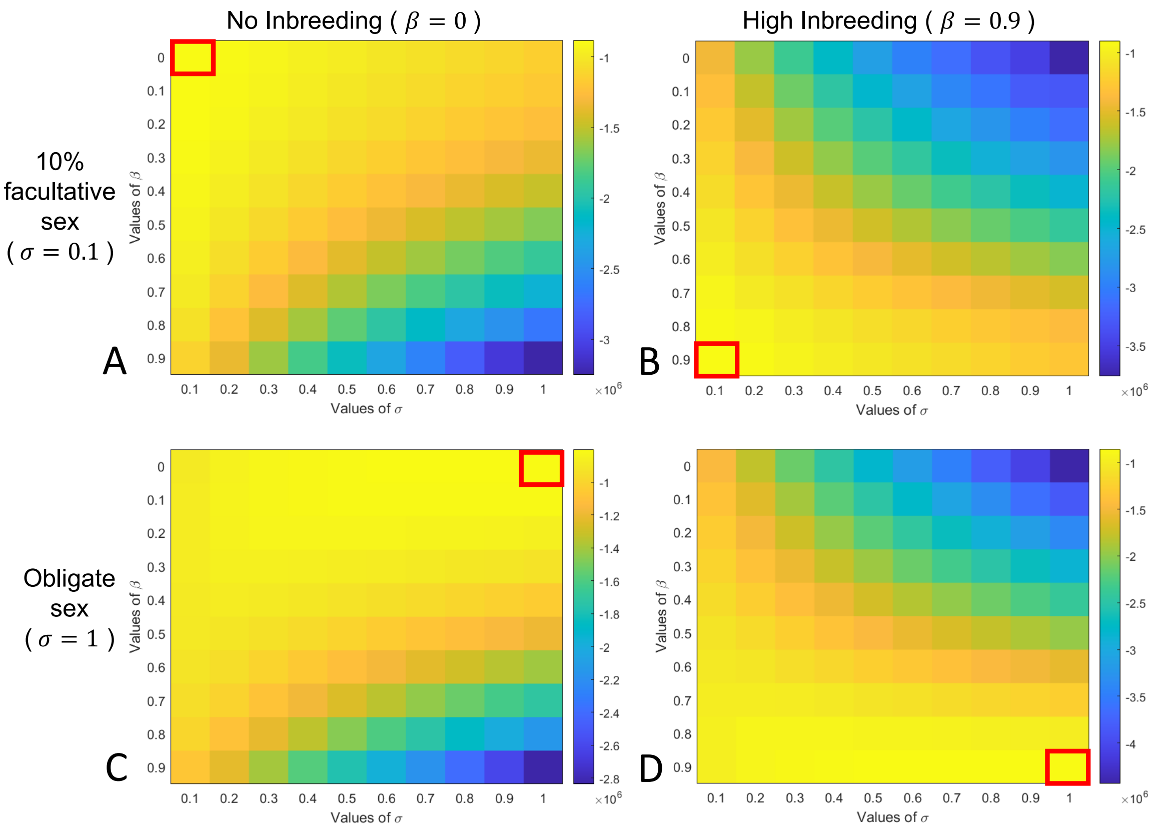


Figure S1: Likelihood surfaces generated from fitting the stationary distribution $P_{\boldsymbol{n}}^{\mathrm{st}}$ of our model to simulated data. The parameters $\beta \in [0, 0.9]$ and $\sigma \in [0.1, 1]$ are varied. The simulated data was sampled every 100 generations and any observations with minor allele frequency appearing less than 5% were filtered out (see Eq. ( S50)). Simulations and models used population size $N = 100$. Model parameters: **A**: $\sigma = 0.1$, $\beta = 0$ (10% facultative sex and no inbreeding); **B**: $\sigma = 0.1$, $\beta = 0.9$ (10% facultative sex and high inbreeding); **C**: $\sigma = 1$, $\beta = 0$ (obligate sex and no inbreeding); **D**: $\sigma = 1$, $\beta = 0.9$ (obligate sex and high inbreeding). The largest log-likelihoods are in yellow and the smallest are in blue. A red box indicates the maximum log likelihood value as the model parameters $\beta$ and $\sigma$ are varied. The parameters that gave the maximum log likelihood of observing each data set are the exact parameters used to simulate the data, when varying parameter values along a grid.

#### S4.2 Fitting the sampled stationary distribution $\mathcal{P}^{\text{st}}_{\boldsymbol{\eta}}$ to simulated data using $N\sigma$ Theory

In the main text (Mathematical Modeling section), we provide mathematical justification for our assumption that two stationary probability distributions, $P^{\text{st}}_{\boldsymbol{n}}(N_1, \sigma_1, \beta)$ and $P^{\text{st}}_{\boldsymbol{n}}(N_2, \sigma_2, \beta)$, will look qualitatively similar if $N_1\sigma_1 = N_2\sigma_2$ (up to the discretisation of the state space). We illustrate this property in the upper panels of Fig. S2. Although the distributions have different population sizes, the distributions look qualitatively similar because the number of sexual events per generation time, $N\sigma$, is conserved across the two populations.

Importantly, when these original stationary distributions, $P^{\text{st}}_{\boldsymbol{n}}(N_1, \sigma_1, \beta)$ and $P^{\text{st}}_{\boldsymbol{n}}(N_2, \sigma_2, \beta)$, are subjected to random sampling to produce the sampled distributions $\mathcal{P}^{\text{st}}_{\boldsymbol{\eta}}(N_s, N_1, \sigma_1, \beta)$ and $\mathcal{P}^{\text{st}}_{\boldsymbol{\eta}}(N_s, N_2, \sigma_2, \beta)$ (see Eq. (7) of main text), the sampled distributions inherit this property. That is, the sampled distributions are approximately the same so long as $N\sigma$ is the same in the two populations (see Fig. S2, lower panels). In the main text (Parameter Fitting section) we describe how we can exploit this property to simplify our parameter fitting algorithm.

Solving the equation for the stationary probability distribution $P^{\text{st}}_{\boldsymbol{n}}(N, \sigma, \beta)$ scales approximately quadratically with $N$ (the state space for $\boldsymbol{n} = (n_{AA}, n_{Aa})$ is of size $(N+1)(N+2)/2$). This makes the distribution $P^{\text{st}}_{\boldsymbol{n}}(N, \sigma, \beta)$ prohibitively expensive to calculate for biologically reasonable $N = N_e$, e.g. $N = \mathcal{O}(10^4)$ (see main text Table 1).

In order to avoid this issue, we *propose* an artificially low value of $N$, $N_{fit}$, to use to obtain a stationary distribution, $P^{\text{st}}_{\boldsymbol{n}}(N_{fit}, \sigma_{fit}, \beta)$ computationally cheaply. We then use this distribution in turn to produce a sampled distribution, $\mathcal{P}^{\text{st}}_{\boldsymbol{\eta}}(N_s, N_{fit}, \sigma_{fit}, \beta)$. We fit this distribution (derived using our artificially low value of $N = N_{fit}$) to our data to obtain an estimate of $\sigma_{fit}$ and $\beta$, $\hat{\sigma}_{fit}$ and $\hat{\beta}$. We can then translate back to the expected value of $\sigma$ in the population data, $\hat{\sigma}$, so long as we have an estimate for the size $N$ of the population from which the data was derived, $N = N_e$; $\hat{\sigma} = N_{fit}\hat{\sigma}_{fit}/N_e$.

To make the above description concrete, suppose that the upper-left panel of Fig. S2 represents the true distribution of a population's genotype frequencies with is $N_e = 200$. We have an estimate for this $N_e$, but not for $\sigma$ and $\beta$ in the population (0.025 and 0.7 respectively). However we do have a sample of $N_s = 40$ individuals in the population, and can use these to construct a sampled distribution (see Fig. S2, lower-left panel). In order to fit the data computationally cheaply, we *propose* $N_{fit} = 100$. We then produce a series of stationary distributions $P^{\text{st}}_{\boldsymbol{n}}(N_{fit}, \sigma_{fit}, \beta)$ (e.g. Fig. S2, upper-right panel) and corresponding sampled distributions (e.g. Fig. S2, lower-right panel). We compare our sampled distributions (e.g. Fig. S2, lower-right panel) to our data sampled from the population (e.g. Fig. S2, lower-left panel), to obtain a maximum liklihood estimate for $\sigma_{fit}$ and $\beta$. In the case of Fig. S2, we would obtain $\hat{\sigma}_{fit} \approx 0.05$ and $\hat{\beta} = 0.7$. In order to obtain our estimate for the population, we take our proposed $N_{fit} = 100$ and multiply this by our maximum likelihood estimate for the rate of sexual reproduction

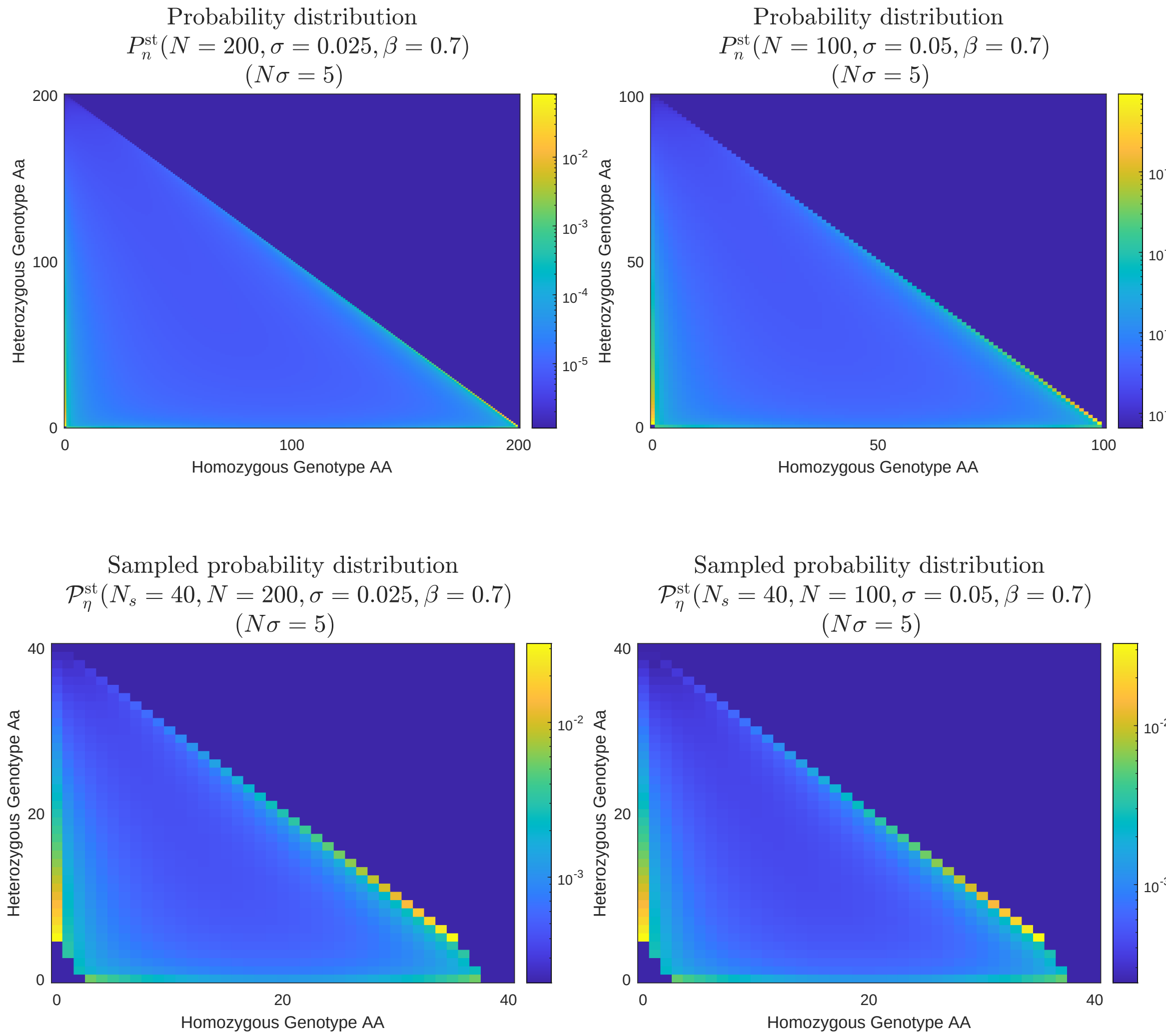


Figure S2: Figure illustrating invariance of distributions while the compound parameter $N\sigma$ is held constant. Upper panels show the stationary probability distribution $P_{\boldsymbol{n}}^{\rm st}$ obtained from solving the master equation for populations of $N = 200$ (top left panel) and $N = 100$ (top right panel), as defined in Eq.(2) of the main text. Although the population size has changed, the distributions remain comparable because the probability of sexual reproduction, $\sigma$ has been increased so that $N\sigma = 5$ in both populations. Lower panels show the sampled stationary distribution $\mathcal{P}_{\boldsymbol{\eta}}^{\rm st}$, as defined in Eq. (7) of the main text.

to obtain an estimate for the number of sexual reproduction events in a generation; $N_{fit}\hat{\sigma}_{fit} = 5$. In order to calculate an estimate for $\hat{\sigma}$ in the population, we then must divide the best fit number of sexual reproduction events in a generation by the true population size $N_e = 200$, to obtain $\hat{\sigma} = 0.025$.

The above represents a somewhat idealised scenario. In the remainder of this section we will use simulated data to explore the effect of changing various fitting parameters in terms of our estimates for $\hat{\sigma}$ and $\hat{\beta}$. In Section S4.2.1 we describe the simulated data. In Section S4.2.2 we describe what happens to the fitting as the parameter $N_{fit}$ is reduced. We see the $N\sigma$ rule emerges empirically from simulated data. However when $N_{fit}$ becomes too small ($50 \gtrsim N_{fit}$), the rate of inbreeding $\hat{\beta}$ is overestimated. This effect stems from the very high level of discretization of the $P_{\boldsymbol{n}}^{\rm st}(N, \sigma, \beta)$ state space when $N$ is small. In Section S4.2.3 we describe the effect of reducing the number of observations of the simulated data (equivalent to a reduced number of segregating sites, $M$ in our data). We find that very low numbers of observations ($1000 \gtrapprox M$) can lead to an over-estimate of $\hat{\sigma}$ and both an over and under-estimate of $\hat{\sigma}$. Finally in Section S4.2.4 we describe what happens to the fitting as the parameter $N_s$ is reduced. We find that a small number of samples ($50 \gtrapprox N_s$) can lead to an underestimate for the rate of sexual reproduction, $\hat{\sigma}$, and overestimate for the rate of inbreeding $\hat{\beta}$.

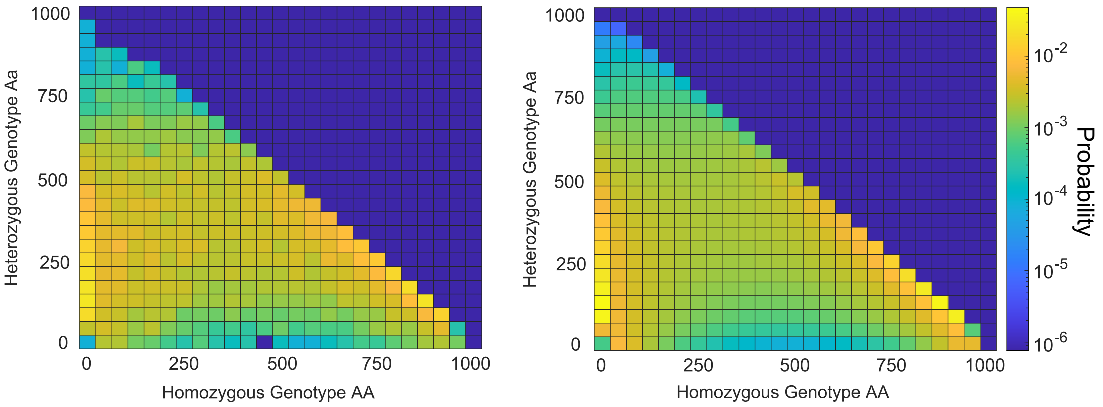


Figure S3: Heatmap of simulated data probability distribution (left) and maximum likelihood stationary probability distribution (right) for a population of $N_{sim} = 1000$. Remaining simulation parameters are $\sigma_{sim} = 0.01$ and $\beta_{sim} = 0.45$. Simulated data was resampled (on average) every 100 generations across approximately $1.3 \times 10^6$ generations (approximately $1.3 \times 10^9$ total birth-death events), and was filtered according to the minor allele frequency condition (see Eq. (S50)). Maximum likelihood parameter estimates: $\hat{\sigma} = 0.01$, $\hat{\beta} = 0.44$. Probability has been binned in both plots, where each bin has height and width 40. Larger probabilities are in yellow and lower probabilities are in blue, using a logarithmic scale.

##### S4.2.1 Simulated Data

To verify the robustness of the model, simulated data was generated using the Gillespie algorithm (stochastic simulation algorithm) and fitted to the model to compare estimated parameter values with corresponding theoretical values. The simulation used the following parameters: $N_{sim} = 1000$, $\sigma_{sim} = 0.01$ ($N_{sim}\sigma_{sim} = 10$), $\beta_{sim} = 0.45$, and initial conditions: $(n_{AA}, n_{Aa}) = (1, 0)$. The total time of the simulation was approximately $T = 1.3 \times 10^6$ generations (approximately $1.3 \times 10^9$ birth-death events), with no burn in time[1]. The data was sampled (on average) every 100 generations, and after minor allele filtering was applied[2] (see Eq. (S50)), a total of 11539 observations remained. The simulated data can be found in [11].

Figure S3 shows a heatmap displaying the probability distribution of the simulated data used for the results in both Figures S4 and S5, and the stationary probability distribution, calculated using the maximum likelihood parameter estimates, $\hat{\sigma} = 0.01$, $\hat{\beta} = 0.44$.

[1] Due to this, observations towards the end of the simulated data set are thought to be closer to equilibrium than those at the beginning.

[2] That is, all observations that satisfy the inequality in Eq. (S50) are retained.

##### S4.2.2 Varying the model fitting population size, $N_{fit}$

Figure S4 displays the results of fitting simulated data to the model using different population sizes, $N_{fit}$, where $N_{fit} \leq N_{sim}$ and $N_{sim}$ is the population size used in the simulation.

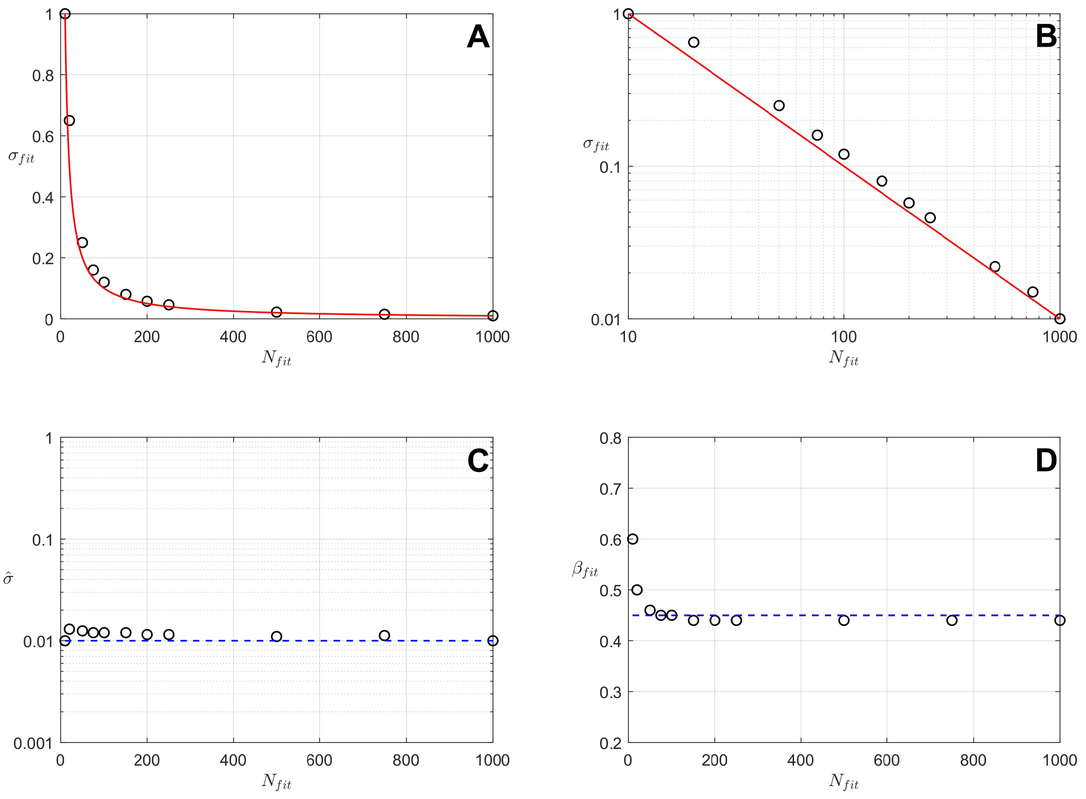


Figure S4: Estimates of model parameters, $\sigma$ and $\beta$, obtained by fitting simulated data observations using different values of the population size, $N_{fit}$. Simulated data generated using simulation parameters: $N_{sim} = 1000$, $\sigma_{sim} = 0.01$ ($N_{sim}\sigma_{sim} = 10$), and $\beta_{sim} = 0.45$. By fitting using a population size $N_{fit} \leq N_{sim}$, the model was able to approximately determine the value of $N_{sim}\sigma_{sim} = 10$ and returned maximum likelihood estimates of $\sigma_{fit}$ which approximately satisfy the relationships described by Eqs. (S53-S54). **A**: Red curve represents the theoretical relationship between $\hat{\sigma}_{fit}$ and $N_{fit}$ (see Eq. (S54)), with observed points shown in black. **B**: Red line represents this same theoretical relationship on a log scale (see Eq. (S55)). **C**: Dashed blue line shows the expected estimate of $\sigma$ in a population of $N_{sim}$ when scaling appropriately (see Eq. (S56)), with observed estimates for different values of $N_{fit}$ in black. **D**: Dashed blue line shows the expected estimate of $\beta$ for different values of $N_{fit}$ with observed estimates in black.

Figure S4**A** and S4**B** shows the maximum likelihood estimates of $\sigma$, labelled as $\hat{\sigma}_{fit}$ for different values of $N_{fit}$ between 10 and 1000, on a both a linear and log scale respectively. The red curve/line represents the theoretical relationship (see Eqs. (S54-S55)), with the estimates $\sigma_{fit}$ in black. As $N_{fit}$

increases, the value of $\hat{\sigma}_{fit}$ decreases, and when $N_{fit} = N_{sim} = 1000$, the value of $\hat{\sigma}_{fit} = \sigma_{sim} = 0.01$:

$$N_{fit}\sigma_{fit} = N_{sim}\sigma_{sim} \tag{S53}$$

$$\sigma_{fit} = \frac{10}{N_{fit}} \tag{S54}$$

$$\log_{10}(\sigma_{fit}) = 1 - \log_{10}(N_{fit}) \tag{S55}$$

such that

$$\hat{\sigma} = \frac{N_{fit}\sigma_{fit}}{N_{sim}}. \tag{S56}$$

Estimates of $\sigma$ for different values of $N_{fit}$ agree with the theoretical relationship, as all observations. Hence the model can correctly identify the value of the compound parameter $N\sigma$, which for the simulated data set was 10.

Rearranging Eq. (S53) and solving for $\sigma_{sim}$ we arrive at Eq. (S56) where $\hat{\sigma}$ represents the maximum likelihood estimate of $\sigma$ in the simulated population of size $N_{sim}$, calculated by fitting a model using a population of size $N_{fit}$ which gave the maximum likelihood estimate $\sigma_{fit}$. Figure S4**C** shows the values of $\hat{\sigma}$ in black with the simulated value of 0.01 in blue, for different values of $N_{fit}$. For all values of $N_{fit}$ the values of $\hat{\sigma}$ lie close to the expected value of 0.01, with slight overestimates for smaller values of $N_{fit}$. Therefore the fitting procedure accurately estimates the value of $\sigma$ in the population of size $N_{sim}$ to within the same order of magnitude, when fitting use a population size $N_{fit}$.

Figure S4**D** shows the maximum likelihood estimates of $\beta$, labelled $\hat{\beta}_{fit}$, for different values of $N_{fit}$, with the expected value of 0.45 in blue. For values of $N_{fit}$ above 50, the values of $\hat{\beta}_{fit}$ agree with the expected value, however for values of $N_{fit}$ below 50, $\beta$ is overestimated. As $N_{fit}$ is a parameter that we are free to choose however, we can enforce $N_{fit} \geq 100$ in our fitting procedure.

##### S4.2.3 Varying the number of observations used in model fitting, $M$

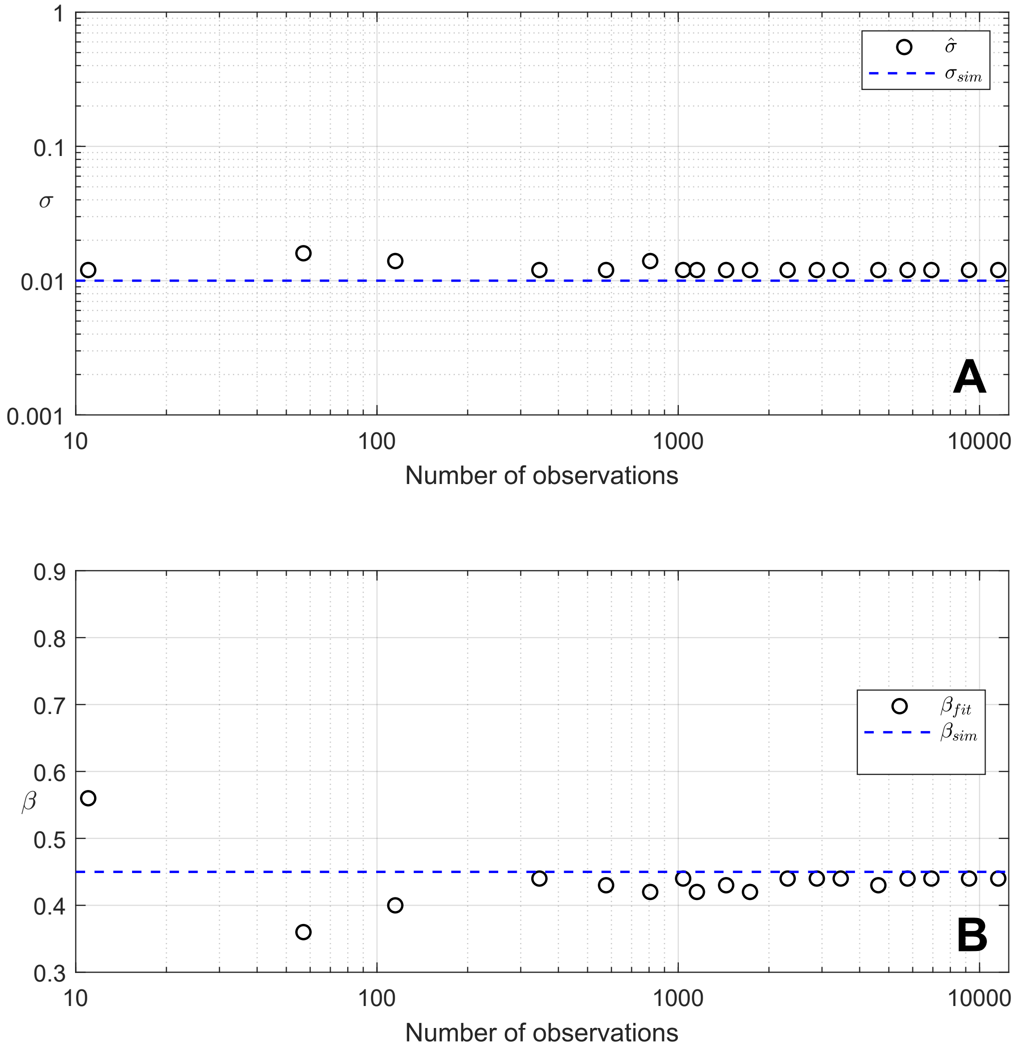


Figure S5: Estimates of model parameters, $\sigma$ and $\beta$, obtained by fitting simulated data observations to a population size of $N_{fit} = 200$, whilst varying the number of observations. Simulated data generated using simulation parameters: $N_{sim} = 1000$, $\sigma_{sim} = 0.01$ ($N_{sim}\sigma_{sim} = 10$), and $\beta_{sim} = 0.45$. Random subsets of the total $11,539$ observations were selected for the model fitting procedure. **A**: Dashed blue line shows the expected estimate of $\sigma$ in a population of $N_{sim}$ when scaling appropriately (see Eq. (S53)) for different numbers of observations, with observed estimates in black. **B**: Dashed blue line shows the simulated value of $\beta$ for different numbers of observations, with estimates of $\hat{\beta}$ in black.

Figure S5 displays the results of fitting different amounts of simulated data to the model using a model of population size $N_{fit} = 200$. The simulated data was the same as that used for the results in Figure S4 (hence used the same simulation parameters), with a total of 11539 observations. Different sized subsets of these observations were randomly selected and the model fitting procedure was applied to these subsets of data. These subsets ranged from 0.1% to 100% of the total observations. Figure S5**A** shows the value of $\hat{\sigma}$ for different numbers of observations, with the expected value of 0.01 shown in blue. As the number of observations decreases below 1000, the values of $\hat{\sigma}$ increase and $\sigma$ is overestimated even further. For the number of observations above 1000, the value of $\hat{\sigma}$ stabilises at the value 0.012, slightly above the expected value of 0.01. Figure S5**B** shows the value of $\hat{\beta}_{fit}$ for different numbers of observations, with the expected value of 0.45 shown in blue. The fewer the observations, the more unreliable the estimates of $\beta$ become. For the number of observations above 1000, estimates of $\beta$ stabilise around 0.44. Since both estimates of $\sigma$ and $\beta$ stabilise after approximately 1000 observations, we note that the results of fitting data sets with less than 1000 observations should be treated with caution.

###### S4.2.4 Varying the number of individuals sampled from the wider population, $N_s$

The data used in this investigation is from a sample of Leishmania parasites from the wider population. To investigate the effect the sample population size has on the estimates of $\sigma$ and $\beta$, samples from the simulated data, of different sample sizes ($N_{samp}$), were randomly selected to create new data sets, of which the model was fitted to. The observations were randomly drawn from a multinomial distribution with replacement, where the sample population size ($N_{samp}$) was fixed for each data set and the probability of drawing each variable was set equal to the genotype frequency observed in the original simulated data. Each new data set was filtered according to Eq. (S50) and was scaled to fit to a population size $N_{fit} = 200$, to estimate $\sigma$ and $\beta$. Subsequently, values of Wright's inbreeding coefficient, $F_{IS}$, were estimated from the data by substituting the maximum likelihood estimate of $\beta$ into Eq. (S57). These inbreeding estimates were compared to estimates using the traditional population approach relevant for obligate sexual reproduction (see Section S2.3).

Figure S6**A** shows the values of $\hat{\sigma}$, the maximum likelihood estimates of $\sigma$ in the population of size $N_{sim} = 1000$, obtained from fitting models to simulated data sampled using different sample population sizes ($N_{samp}$). As the proportion of the simulated population sampled decreases from 100% sampled, to 1% sampled, the value of $\hat{\sigma}$ decreases from approximately the expected theoretical value of 0.01, to approximately 0.003. As the percentage of the population sampled decreases below 10%, $\sigma$ is underestimated. This may indicate that for the data sets EA2 and ISC2, which had sample population sizes 18 and 15 respectively, the values of $\hat{\sigma}_e$ may be underestimates.

Figure S6**B** displays the estimates of $\beta$ and $F_{IS}$ obtained from traditional population genetics calculations that assume high rates of sexual reproduction (see Section S2.3) and from fitting models to simulated data sampled using different sample population sizes ($N_{samp}$). Figure S6**B** shows the estimates of $\beta$, with model estimates in black, population genetics estimates in red, and the theoretical value of 0.45 in blue. Similarly, Figure S6**B** shows the corresponding estimates of $F_{IS}$. The model estimates of $\beta$ and $F_{IS}$ align with the theoretical value of 0.45 and 0.29 respectively, whereas when using the population genetics calculation, $\beta$ and $F_{IS}$ are consistently underestimated, approximately 0.25 and 0.14 respectively. For the model estimates of $\beta$ and $F_{IS}$, as the proportion of the simulated population sampled decreases from 100% sampled, to 1% sampled, the values of $\beta$ and $F_{IS}$ increase above the expected theoretical values. As the percentage of the population sampled decreases below 4%, $\beta$ and $F_{IS}$ are overestimated.

This may indicate that for the data sets EA2 and ISC2, which had a small number of individuals sampled, 18 and 15 respectively, the values of $\hat{\beta}$ and $\hat{F}_{IS}$ may be overestimates.

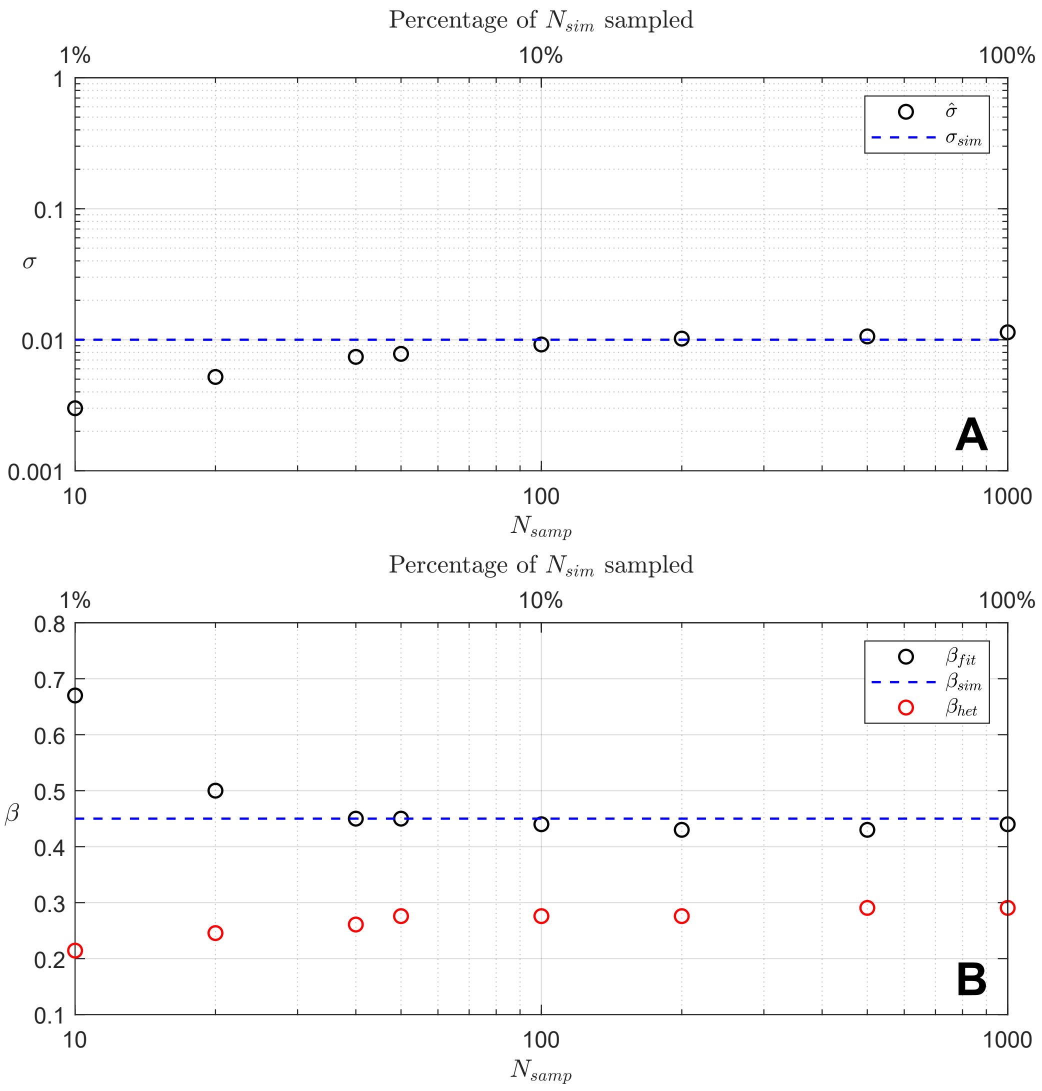


Figure S6: Estimates of $\sigma$ and $\beta$ obtained by sampling each observation from the simulated data (see Section S4.2.1) using different sample sizes ($N_{samp}$) and fitting the sampled data to models of size $N_{fit} = 200$. Random samples for each observation were drawn from a multinomial distribution with replacement, with the probability of drawing each variable $(n_{AA}, n_{Aa})$ equal to the genotype frequency at each observation. Simulation parameters: $N_{sim} = 1000$, $\sigma_{sim} = 0.01$, $\beta_{sim} = 0.45$. **A**: Shows estimates of $\sigma$ with model estimates ($\hat{\sigma}$) in black, and the simulation value ($\sigma_{sim} = 0.01$) in blue. **B**: Shows estimates of $\beta$ with model estimates in black ($\beta_{fit}$), estimates calculated from heterozygosity values ($\beta_{het}$) in red, and the simulation value ($\beta_{sim} = 0.45$) in blue. Estimates of $\beta_{het}$ were inferred from Wright's inbreeding coefficient, as detailed in Section S5.2. Estimates are plotted against the percentage of the simulated population that was sampled.

### S5 Importance of co-fitting inbreeding rate and rate of sexual reproduction

#### S5.1 Estimation of Wright's inbreeding coefficient using $\beta$

The rate of inbreeding, $\beta$, can be converted into Wright's inbreeding coefficient ($F_{IS}$) using the following formula [7]:

$$F_{IS}(\beta) = \frac{\beta}{2-\beta} \tag{S57}$$

which was used to obtain the estimates $F_{IS}(\beta)$ in Table 1 in the main text. Similarly Eq. (S57) can be rearranged to express $\beta$ in terms of $F_{IS}$:

$$\beta = \frac{2F_{IS}}{1+F_{IS}}. \tag{S58}$$

#### S5.2 Estimation of Wright's Inbreeding coefficient using expected and observed heterozygosity

| **Data set** | $F_{IS}$ | $\beta_{Het}$ |
|---|---|---|
| EA1 | 0.133 | 0.235 |
| EA2 | 0.688 | 0.816 |
| ISC1 | 0.492 | 0.659 |
| ISC2 | 0.633 | 0.776 |
| Brazil | −0.144 | −0.337 |

Table S3: Estimates of Wright's inbreeding coefficient ($F_{IS}$) using the heterozygosity data (see Eq. (S57)), and corresponding values of $\beta$ (see Eq. (S58)) for each Leishmania data set.

Estimates of Wright's inbreeding coefficient, $F_{IS}$, were calculated using the traditional population genetics approach to compare those estimates to ones from the model. These estimates are independent of the estimates obtained using $\beta$ in Eq. (S57). This estimate involved calculating the average expected heterozygosity and the average observed heterozygosity across all sites (loci) in the sample, and substituting these into the following equation [6]:

$$F_{IS} = \frac{\langle H_{exp}\rangle - \langle H_{obs}\rangle}{\langle H_{exp}\rangle} \tag{S59}$$

where $\langle H_{exp}\rangle$ is the average expected heterozygosity ($\langle H_{exp}\rangle = H_{exp}/M$, where $M$ is the total number of sites), and $\langle H_{obs}\rangle$ is the average observed heterozygosity ($\langle H_{obs}\rangle = H_{obs}/M$ where $M$ is the total number of sites). The average observed heterozygosity is calculated by simply averaging the observed heterozygosity across all sites in the sample. To calculate the expected heterozygosity, the population is assumed to be at the Hardy-Weinberg equilibrium, so that we have

$$H_{exp} = 1 - p^2 + q^2 \tag{S60}$$

where $p^2$ and $q^2$ are the expected homozygosities of $AA$ and $aa$ genotypes respectively, and $p$ and $q$ are the allele frequencies of $A$ and $a$. The allele frequencies can be calculated from the data at each site, such that:

$$p = \frac{2n_{AA} + n_{Aa}}{2N} \tag{S61}$$

$$q = 1 - p. \tag{S62}$$

Substituting these values into Eq. (S60) and averaging over the number of sites returns the average expected heterozygosity.

Alternatively, an estimate of $F_{IS}$ can be calculated using the aforementioned method for each individual in the sample, and can then be averaged across all individuals to return a single sample estimate. These estimates were almost identical to estimates obtained using Eqs. (S59-S62), so are omitted for succinctness.

Table S3 shows estimates of $F_{IS}$ obtained from each Leishmania data set using Eq. (S59) and the corresponding estimate of $\beta$ (denoted as $\beta_{Het}$), calculated by substituting these values of $F_{IS}$ into Eq. (S58).

#### S5.3 Estimation of the rate of facultative sexual reproduction in the effective population $\sigma_e$

The rate of facultative sexual reproduction in the effective population $\sigma_e$ can be calculated from an estimate of $\sigma$ obtained from fitting data to a model using a smaller population size than $\hat{N}_e$ ($N_{fit} \ll \hat{N}_e$) and an estimate of the effective population size. Since the model can distinguish between different values of the compound parameter $N\sigma$, the following relationship is proposed:

$$N_{fit}\sigma_{fit} = \hat{N}_e\hat{\sigma}_e \tag{S63}$$

assuming that $\sigma_{fit}$ is of order $1/N_{fit}$. Rearranging Eq (S63) to solve for $\hat{\sigma}_e$ gives:

$$\hat{\sigma}_e = \frac{N_{fit}\sigma_{fit}}{\hat{N}_e} \tag{S64}$$

which, after substituting in the values of $N_{fit}$, $\sigma_{fit}$, and $\hat{N}_e$, returns the estimates of $\hat{\sigma}_e$ in Table 1 in the main text. See Section S2.4 for mathematical justification of this relationship.

### S6 Results of model fitting

This sections contains visualisations of the Likelihood surfaces, genotype data and best-fit stationary probability distributions for the various *Leishmania* samples used in this paper:

- EA1 (see Fig. S7)
- EA2 (see Fig. S8)
- ISC1 (see Fig. S9)
- ISC2 (see Fig. S10)
- Brazil (see Fig. S11) .

## S6.1 EA1

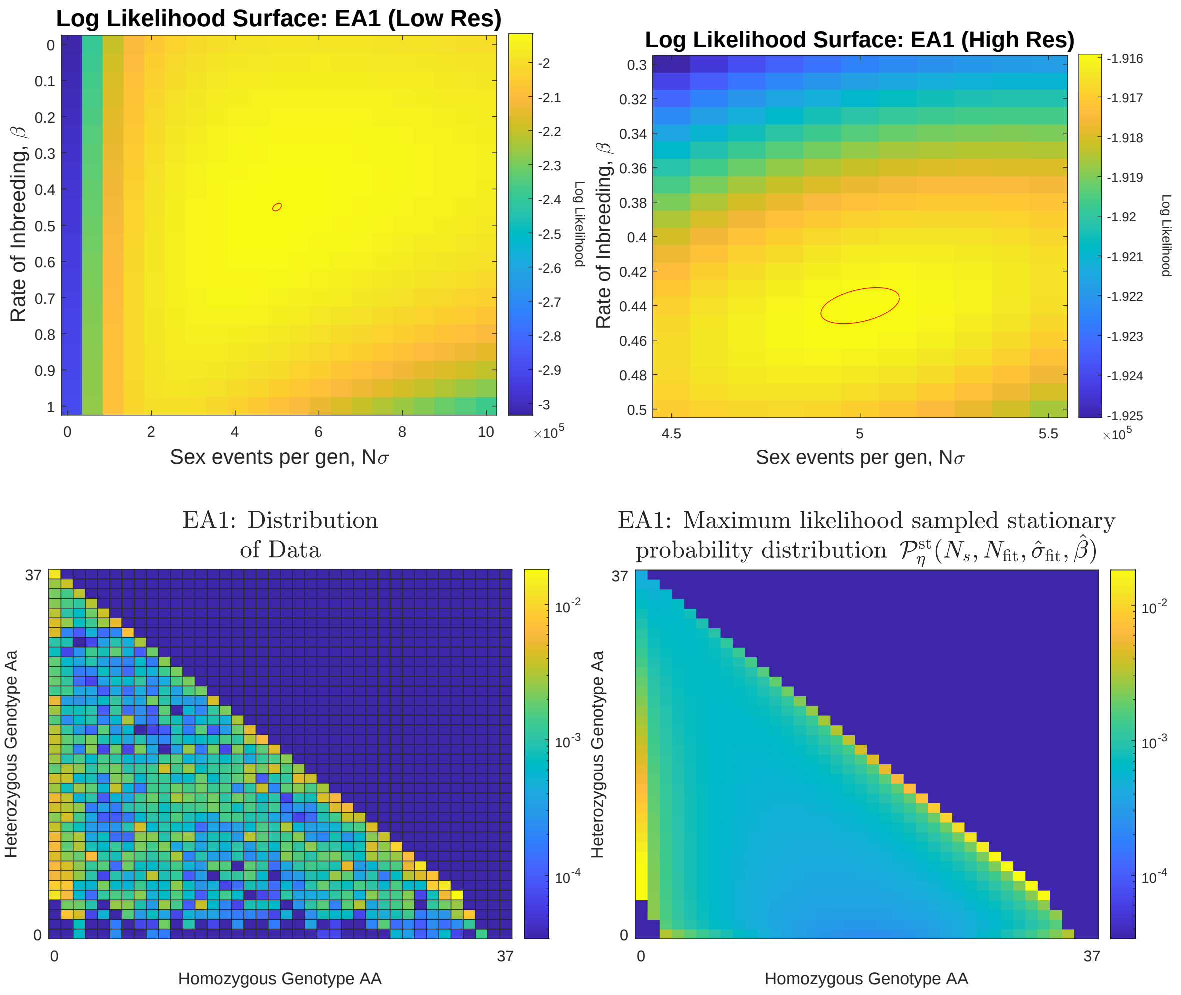


Figure S7: EA1 Likelihood surfaces, data and best fit model distributions. Top panels: Likelihood surfaces in the $N\sigma$-$\beta$ plane. Red circles indicate a 95% confidence interval based for the peak of the likelihood surface based on a covariance matrix derived from Fisher Information matrix (see Statistical analysis in the main text and note that this is an underestimate on the uncertainty as it assumes unlinked loci). Bottom panels: The genotype data (left) and the best-fit sampled stationary probability distribution $\mathcal{P}^{\text{st}}_{\boldsymbol{\eta}}(N_s, N_fit, \hat{\sigma}_{fit}, \hat{\beta})$ (right).

## S6.2 EA2

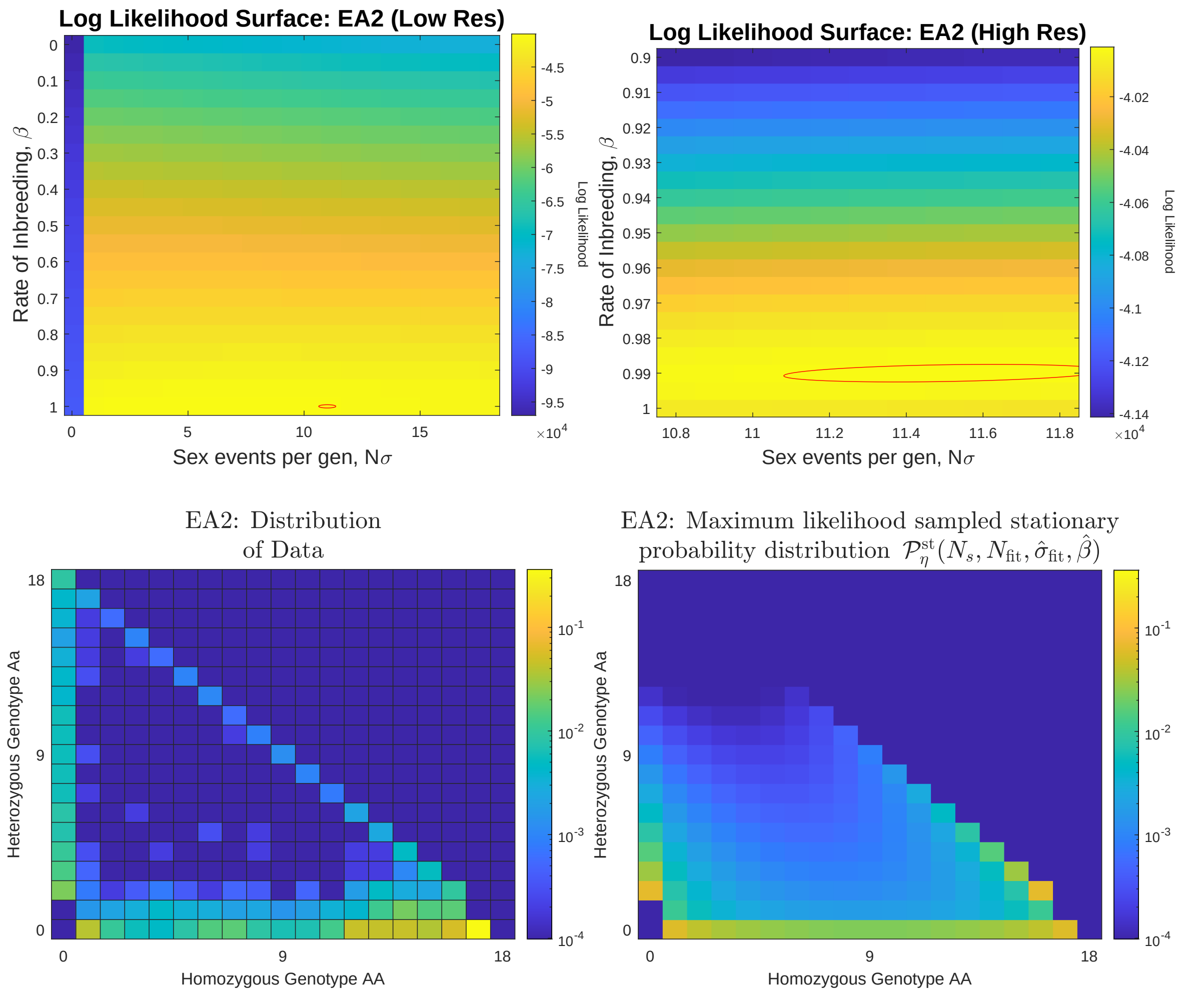


Figure S8: EA2 Likelihood surfaces, data and best fit model distributions. Top panels: Likelihood surfaces in the $N\sigma$-$\beta$ plane. Red circles indicate a 95% confidence interval based for the peak of the likelihood surface based on a covariance matrix derived from Fisher Information matrix (see Statistical analysis in the main text and note that this is an underestimate on the uncertainty as it assumes unlinked loci). Bottom panels: The genotype data (left) and the best-fit sampled stationary probability distribution $\mathcal{P}^{\rm st}_{\boldsymbol{\eta}}(N_s, N_f it, \hat{\sigma}_{fit}, \hat{\beta})$ (right).

#### S6.3 ISC1

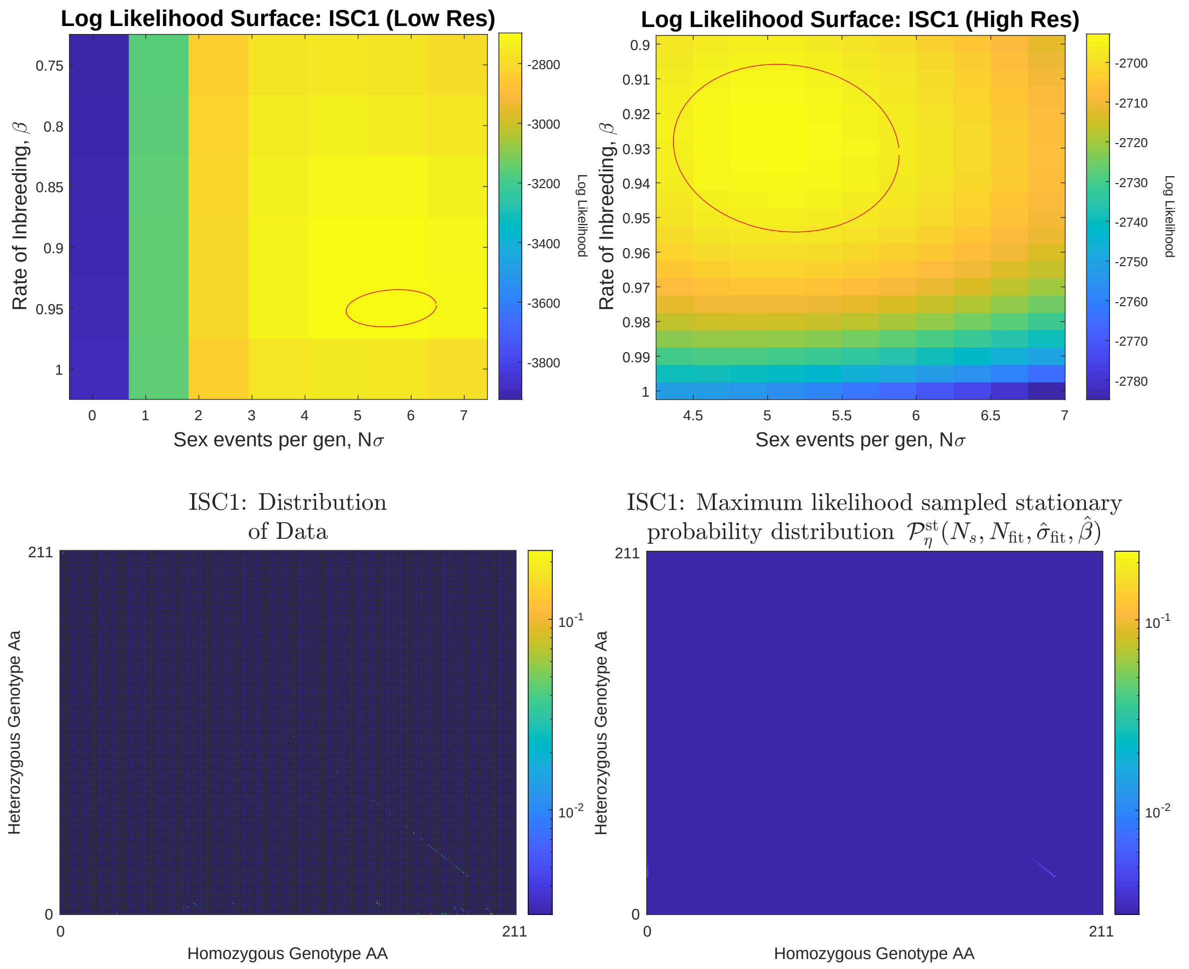


Figure S9: ISC1 Likelihood surfaces, data and best fit model distributions. Top panels: Likelihood surfaces in the $N\sigma$-$\beta$ plane. Red circles indicate a 95% confidence interval based for the peak of the likelihood surface based on a covariance matrix derived from Fisher Information matrix (see Statistical analysis in the main text and note that this is an underestimate on the uncertainty as it assumes unlinked loci). Bottom panels: The genotype data (left) and the best-fit sampled stationary probability distribution $\mathcal{P}^{\rm st}_{\boldsymbol{\eta}}(N_s, Nfit, \hat{\sigma}_{fit}, \hat{\beta})$ (right). Note that the small number of segregating sites in this data set, combined with the large sample size (see Table 1) make the genotype plots particularly difficult to visualise.

#### S6.4 ISC2

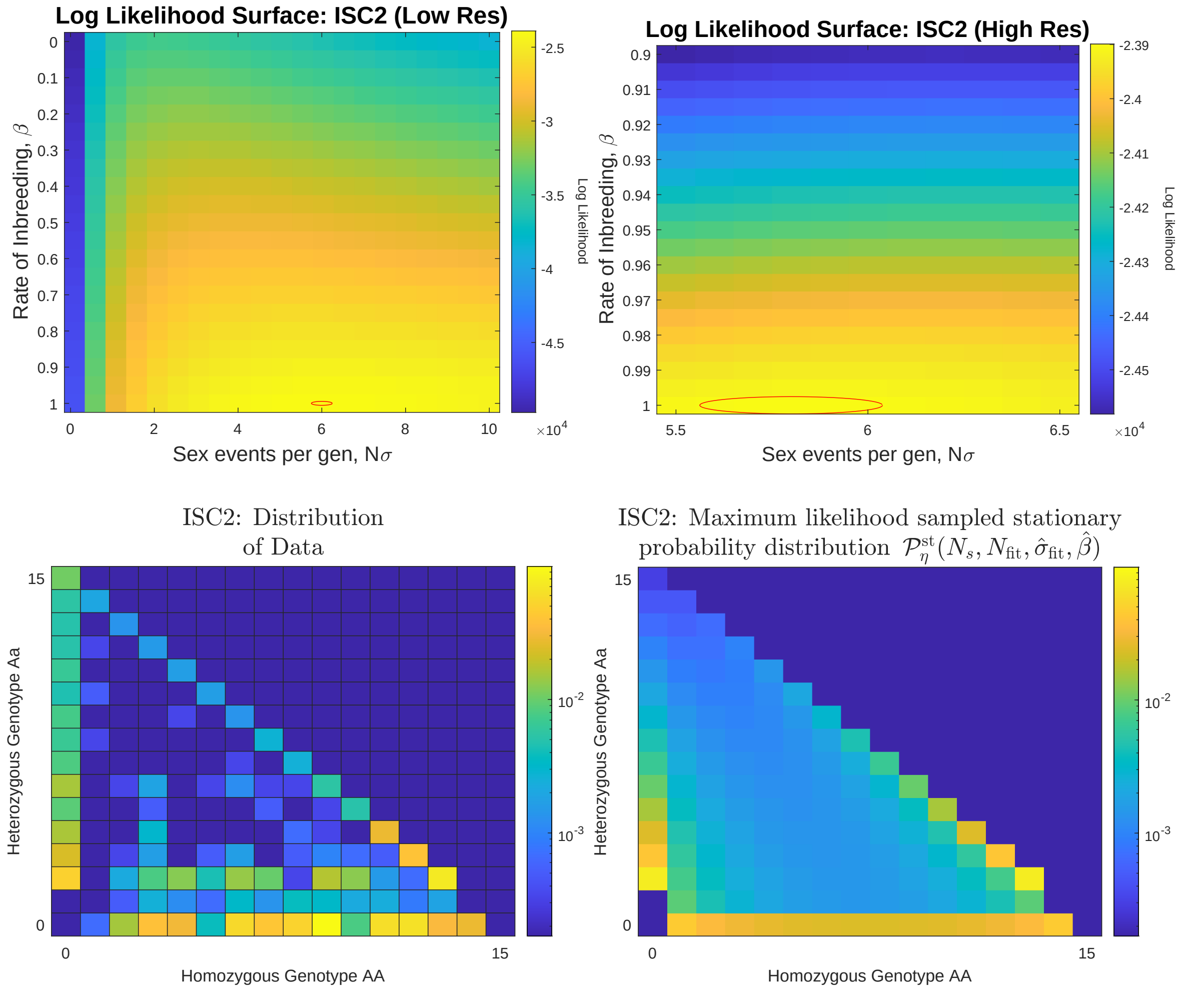


Figure S10: ISC2 Likelihood surfaces, data and best fit model distributions. Top panels: Likelihood surfaces in the $N\sigma$-$\beta$ plane. Red circles indicate a 95% confidence interval based for the peak of the likelihood surface based on a covariance matrix derived from Fisher Information matrix (see Statistical analysis in the main text and note that this is an underestimate on the uncertainty as it assumes unlinked loci). Bottom panels: The genotype data (left) and the best-fit sampled stationary probability distribution $\mathcal{P}^{st}_{\boldsymbol{\eta}}(N_s, N_fit, \hat{\sigma}_{fit}, \hat{\beta})$ (right).

#### S6.5 Brazil

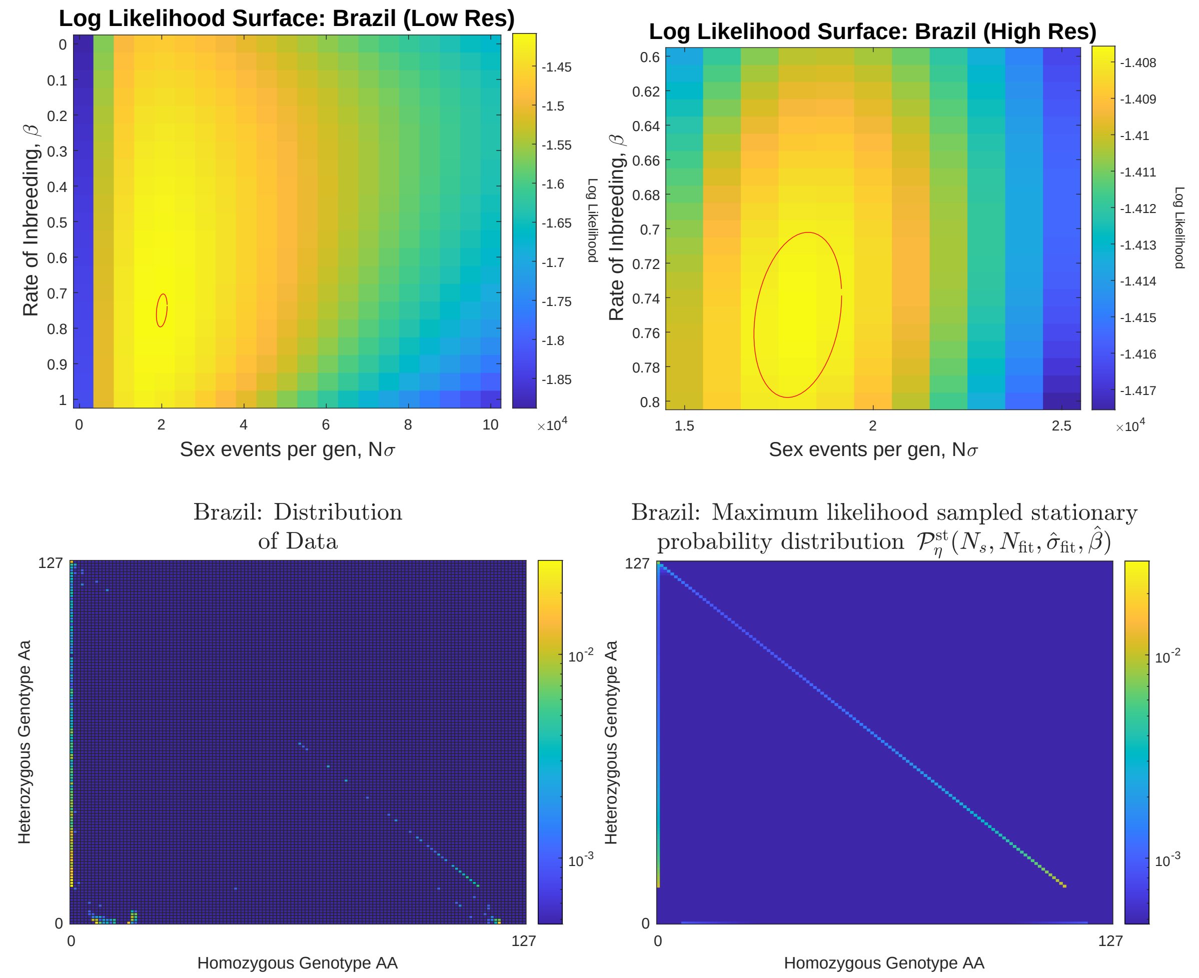


Figure S11: Brazil Likelihood surfaces, data and best fit model distributions. Top panels: Likelihood surfaces in the $N\sigma$-$\beta$ plane. Red circles indicate a 95% confidence interval based for the peak of the likelihood surface based on a covariance matrix derived from Fisher Information matrix (see Statistical analysis in the main text and note that this is an underestimate on the uncertainty as it assumes unlinked loci). Bottom panels: The genotype data (left) and the best-fit sampled stationary probability distribution $\mathcal{P}^{\text{st}}_{\boldsymbol{\eta}}(N_s, N_f it, \hat{\sigma}_{fit}, \hat{\beta})$ (right).

### S7 de Finetti Diagrams

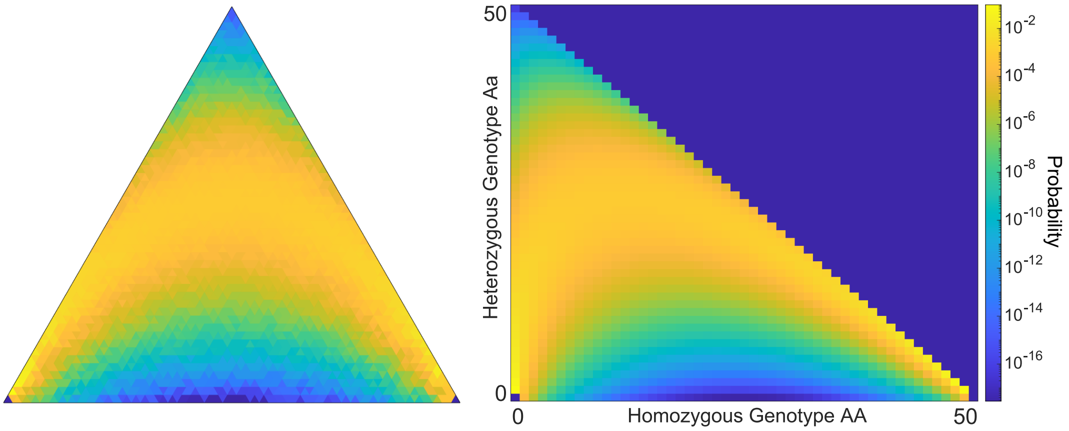


Figure S12: Comparison of the stationary probability distribution as a ternary plot heatmap (left) and normal plot heatmap (right) for model parameters: $N = 50$, $\sigma = 1.0$, $\beta = 0.0$. Ternary plot is the same as that in the centre top panel of Figure 2 in the main text. The highest probabilities are shown in yellow, with the lowest shown in blue, using a log scale. Symmetry of the distribution is visually preserved in the right plot, but not in the left plot due to a discrepancy between the number of states in the state space and the number of triangles that constitute the ternary plot.

The number of triangles that make up the ternary plot (de Finetti diagram) follows the sequence of square numbers. The number of states in the state space follows the sequence $(N + 1)(N + 2)/2$, where $N$ is the population size. Ideally each smaller triangle in the ternary plot would represent a unique state in the state space, however since these sequences do not intersect for any value of $N \in \mathbb{N}$, there exists no population size $N$ such that the number of triangles equals the number of states. This leaves us with the visual discrepancy seen in Figure S12 as well as Figures 2 and 3 in the main text - that the de Finetti diagrams are not exactly symmetric. Despite this, symmetry was preserved in all calculations involving the model, therefore are results are not affected by this issue.

### References


[1] Crispin Gardiner. *Stochastic Methods.* Springer, 2009.
